# Age-dependent changes in mean and variance of gene expression across tissues in a twin cohort

**DOI:** 10.1101/063883

**Authors:** Ana Viñuela, Andrew A Brown, Alfonso Buil, Pei-Chien Tsai, Matthew N Davies, Jordana T Bell, Emmanouil T Dermitzakis, Timothy D Spector, Kerrin S Small

## Abstract

Gene expression changes with age have consequences for healthy aging and disease development. Here we investigate age-related changes in gene expression measured by RNA-seq in four tissues and the interplay between genotypes and age-related changes in expression. Using concurrently measured methylation array data from fat we also investigate the relationship between methylation, gene expression and age. We identified age-dependent changes in mean levels of gene expression in 5,631 genes and in splicing of 904 genes. Age related changes were widely shared across tissues, with up to 60% of age-related changes in expression and 47% on splicing in multi-exonic genes shared; amongst these we highlight effects on genes involved in diseases such as Alzheimer and cancer. We identified 137 genes with age-related changes in variance and 42 genes with age-dependent discordance between genetically identical individuals; implying the latter are driven by environmental effects. We also give four examples where genetic control of expression is affected by the aging process. Analysis of methylation observed a widespread and stronger effect of age on methylation than expression; however we did not find a strong relationship between age-related changes in both expression and methylation. In summary, we quantified aging affects in splicing, level and variance of gene expression, and show that these processes can be both environmentally and genetically influenced.

## Background

Aging is a complex process, characterized by a progressive decline in an organism's biological function and phenotypic characteristics, which leads to an increased chance of developing disease and ultimately the death of the organism [1]. Others have attempted to understand the aging process by identifying common denominators of aging in different organisms [2]. Many of these hallmarks, such as genome instability, epigenetic alterations, loss of proteostasis and telomere attrition, are accompanied by changes in gene expression. Identification of genes differentially expressed with age has proven useful in identifying pathways whose behavior is modified by age, as well as identifying biomarkers of aging and therapeutic targets [3–5]. Expression studies into aging using animal models have discovered that the expression of up to 75% of genes are associated to aging [6]. As well as associations with the level of gene expression, age-related differences in the splicing of the mRNA produced and differences in the genetic regulation of gene expression have also been observed [6, 7]. On the other hand, human studies have only recently managed to identify thousands of genes associated with age in multiple tissues [3, 8, 9], but are still far from identifying the same scale of aging effects in expression or the same variety of changes. Reasons for this include a reduced power to see interactions due to inability to control environment compared to model organisms, and importantly the lack of sufficient human expression data using appropriate technologies and tissues.

In this study, we investigate changes in gene expression with age using previously published RNA-seq measurements of fat, skin, whole blood and derived lymphoblastoid cell lines (LCLs) expression from ∼800 monozygous (MZ) and dizygous (DZ) adult female twins (Additional Table S1). We take a comprehensive approach that includes not only an analysis of the effect of age on the mean of gene expression and alternative splicing, but also look at age-related changes in gene expression variance and changes in genetic regulation. Although age-related changes in variance of gene expression have been identified before [10–14], we believe this is one of the first studies exploring the underlying causes for changes in variance with age. For that and exploiting the similarities and differences between twins, we discover genes where the discordance between MZ pairs is age-dependent implying environmentally driven effects. Also partition of the variance shows that for a majority of genes the effect of age on expression is small relative to the effect of genetic variation. For a more mechanistic understanding of how age affects expression, we studied age-related epigenetic changes using genome-wide methylation profiles from the same fat biopsies as the expression data which showed a more widespread and stronger influence of aging. Finally, we observe a greater degree of sharing of age effects across tissues than has been previously reported, demonstrating the benefits of studying expression using accurate RNA-seq derived quantifications in a large cohort.

## Results

### Effects of aging in gene expression levels

To investigate the wide range of changes in gene expression with age, we used publicly available RNA-seq data from 855 healthy individuals drawn from the TwinsUK cohort ([15, 16], Additional Table S1) in four tissues: i) photo protected skin, ii) subcutaneous fat, iii) whole blood and iv) lymphoblastoid cell lines (LCLs). We consider a gene associated with age if at least one exon was associated with chronological age. We discovered that 36.6% of tested genes (5,631 of 15,353) had expression levels of at least one exon where expression level was significantly associated with age in at least one tissue (FDR < 0.05; Figure 1A, Additional Table S2 and S3, Additional File 1). This number is roughly double that we previously reported (18.3%, 3,019 genes) using exactly the same skin, fat and LCLs samples but measuring expression using microarrays [3] (Figure 1A, Additional Figure S1). This increase in power is due to the higher resolution provided by RNA-seq data, which also allows exon level quantifications which are more biologically interpretable than working on the gene level [17]. We also found that the total number of expressed genes increased as a function of age in fat tissue (adjusted *P* value = 0.00264, Additional Figure S2) but not in the other tissues. Application of Gene Set Enrichment Analysis (GSEA) to the differentially expressed genes showed significant enrichment in GO terms (FDR < 0.05) related to RNA processing, fat metabolism and oxidation reduction in skin; and cell adhesion, membrane structure and sodium channel complex structure in fat tissue (Additional files 2). In blood, there was no specific enrichment for GO terms, and in LCLs only 7 genes showed significant association with age, 3 of which were previously reported [3]. In conclusion, we identify thousands of genes whose expression levels were associated to age.

**Figure 1.**
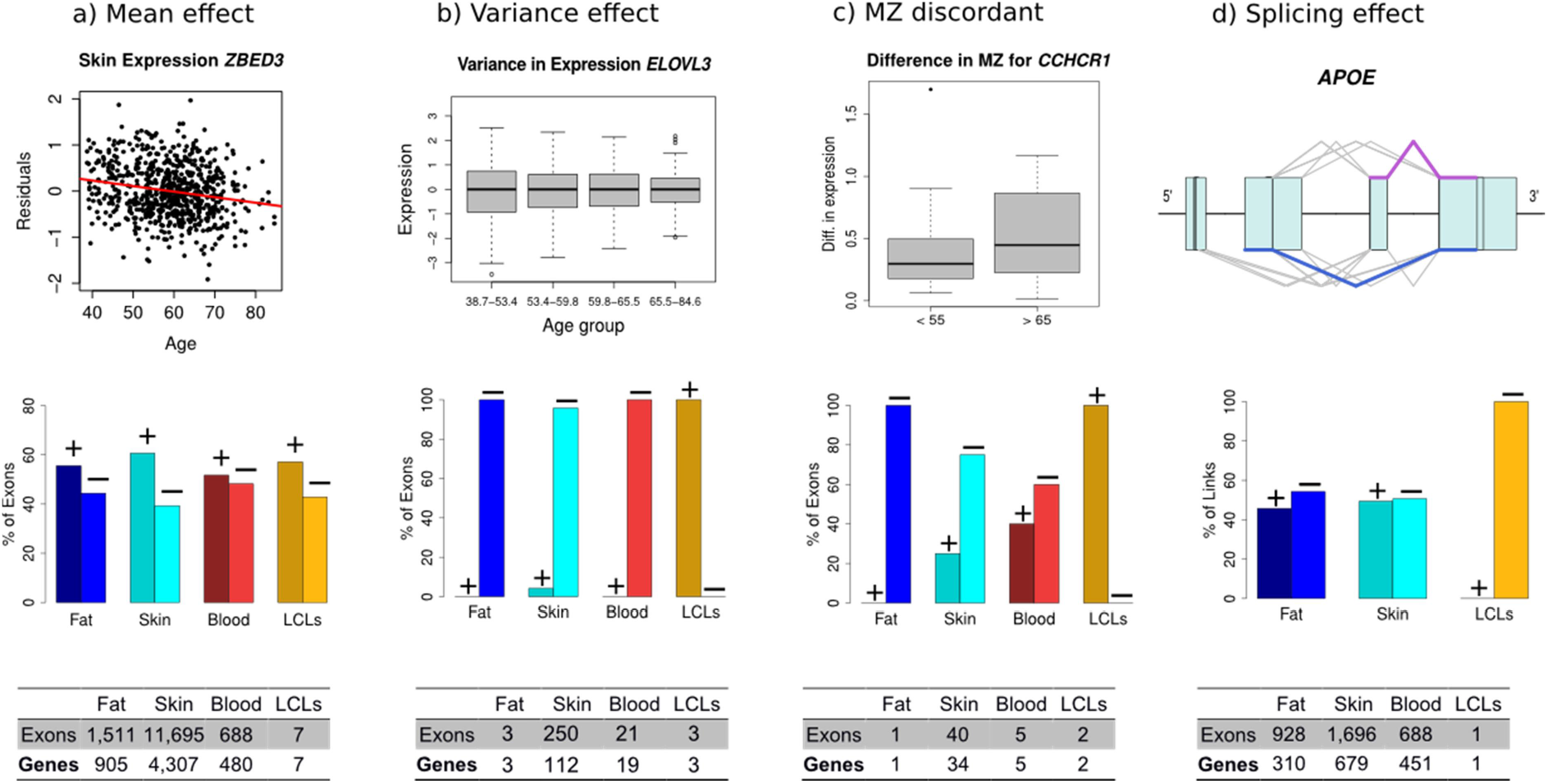
Effects of aging in gene expression: The effect of aging in gene expression is not limited to changes in mean expression values with age (a), but includes also changes in levels of phenotypic variance(b,c), and splicing (d). The top row graphs show real data examples for the effects of aging in expression investigated. The middle graphs show bar plots with the percentage of exons with positive (+) or negative (−) age effects in each analysis. And finally the bottom tables provide the number of exons and genes with significant association for each of the effects presented. All the real examples are from skin, the tissue with larger age effect in expression overall analyses. a) Effect of aging in mean gene expression, usually referred as differentially expression with age in exons. The example shows the residuals (after removing technical covariates) of the expression of the *ZBED3* gene decreasing with age. Skin is the tissues with a larger effect of age in expression and LCLs the smaller. b) The effect of aging in variance of gene expression is shown with the *ELOVL3* gene and a significant decrease of variance in expression with age. From the bar plot it is possible to appreciate that the majority of the significant exons had a decrease in variance with age. c) Differences in expression between monozygous (MZ) twins point out to environmental factors different among the siblings affecting gene expression, since MZ twins are genetically identical individuals with the same age. The example shows the difference in expression between MZ twins in the gene *CCHR1*. d) For the splicing analysis, only links (reads between two exons) were considered. The example shows the structure of the gene *APOE* with its exons (boxes) and lines connecting the exons representing reads spanning between two exons. The number of reads linking exons 3 and 4 (in purple) decreased in number with age, while reads linking exons 2 and 4 (blue) increased with age. The model suggested that an isoform skipping the third exon (from the 5') may be more abundant in older individuals compare to an isoform that includes the third exon linked to the last exon.

To quantify the overall influence of age on global gene expression, we estimated the proportion of variance in exon expression levels (removing technical confounders) explained by age and additive genetic effects (heritability). In exons associated with age, age explained only a small proportion of the variation in gene expression, with median values between 2.2% and 5.7% depending on tissue and with maximum values ranging from 12% to 27% (Additional File 3). Globally, the effect of age on expression was greatest in blood, then skin, fat, and finally LCLs had the least. In comparison, the proportion of variance explained by additive genetic effects on the same set of age-affected exons was greater than that explained by age in all tissues (median *h*^2^_skin_ = 0.12, *h*^2^_fat_ = 0.22, *h*^2^_LCLs_ = 0.20, *h*^2^_blood_ = 0.23). Our results indicate that the influence of age in gene expression is much lower in comparison with the effects of genetic variation in the expression of genes. These relative differences suggest that considering genetics would be crucial in producing an expression derived individual level estimate of “biological age”, similar to those previously proposed for methylation [18].

**Table 1:** Summary of median proportion of variance attributed to age, genetics, common environment and unique environment, for all exons (All) and for age-affected exons (Age). In general, age explained a small proportion of the variance attributed to gene expression. However, for exons affected by age in their expression, the genetic component (heritability) explained significantly higher proportion of the variance in expression compare to the rest of the genes in fat, skin and blood tissues (willconox test Pvalue < 2.1e-17).

|  | Fat |  | Skin |  | Blood |  | LCLs |  |
| --- | --- | --- | --- | --- | --- | --- | --- | --- |
|  | All | Age | All | Age | All | Age | All | Age |
| **Age** | 0.0012 | 0.0287 | 0.0026 | 0.0224 | 0.0039 | 0.0542 | 0.0006 | 0.0366 |
| **Genetics** | 0.0809 | 0.2228 | 0.0856 | 0.1275 | 0.1301 | 0.2333 | 0.1089 | 0.2032 |
| **Commo Env.** | 0.0236 | 0.0573 | 0.0000 | 0.0162 | 0.0380 | 0.0341 | 0.0971 | 0.1312 |
| **Unique Env.** | 0.8550 | 0.6655 | 0.8566 | 0.7662 | 0.7403 | 0.6214 | 0.7320 | 0.6214 |
| **N. of exons** | 101,133 | 1,511 | 96,736 | 11,695 | 71,393 | 688 | 98,372 | 7 |

### Splicing is associated with aging

As well as changes in the average level of expression, age has been shown to cause genome wide changes in splicing [19–21], including in the human brain, that manifest in a differential expression of isoforms with age. Furthermore, splicing changes have been shown to be crucial in the development of a number of age related diseases. To identify changes in splicing with age, we quantified splicing based on link reads between exons using Altrans [22]. This method uses the proportion of reads linking different exons as a quantitative phenotype for splicing in our population, which can be tested for associations with age. We found a total of 904 genes (6.3% of the 14,261 genes with more than one exon expressed) with at least one link differentially expressed with age in either fat or skin (FDR < 0.05, Additional Table S4, Additional Files 4, 5 and 6). For 51.8% of those genes in skin and 11.4% in fat, age was associated with level of expression of the gene as well. We did not see significant associations between age and splicing in LCLs and blood, probably due to the smaller age effect in LCL expression and the smaller sample size available for blood (N = 384). Among genes with age-related differential splicing we found *APOE* (Figures 1D and 5)(previously associated to extreme longevity, Alzheimer disease and cholesterol metabolism), *LMNA* (causal of progeria, an accelerated aging syndrome), *HTRA2* (Parkinson disease) and *AAP* (Alzheimer disease). In fat tissue, many thrombospondins and collagen genes had age-related changes in splicing, as well as genes such as *AKT1* and *AKT2* from the insulin-IGF1 pathway, which is known to play a central role in aging. Overall, we observed that some age-related changes in gene expression were associated to changes in splicing, but that the effect of age in splicing was weaker than the observed effect on expression levels.

### Variance and differences in gene expression between MZ twins is dependent of age

Age-dependent changes in the variance of gene expression (rather than mean expression levels) have been reported in model organisms [10–12] and humans [13, 14]. Changes in phenotypic variance with age can be due to different responses to environment, age-related damage accumulation leading to stochastic deregulation of gene expression or gene-age interactions, where changes in relative genetic effects can increase heterogeneity across the population at a particular age [23]. We looked for changes in variance with age and identified 137 genes where expression showed age-dependent variance in at least one tissue (FDR < 0.05, Figure 1B, Additional File 7). Since changes in global phenotypic variance have mainly been reported to increase with age, we were surprised to observe that for the majority of these genes we report a decrease in variance of expression. This reduction in variance could reflect a decreased ability for expression levels to respond to environmental stimuli in older individuals, or a reduction in genetic regulation with age. The biological functions associated to the genes with age-associated differential variance in skin included oxidation reduction, with examples such as *SOD2*, fatty acid metabolism with genes including *CPT1B*, *ELOV3* and *ELOV5* or cell cycle control like *p21* (Figure 1B, Additional Figure S3). In blood, enriched pathways included the VEGF signaling pathway with the *PIK3CD* and *PXN* genes. Our analysis shows concrete examples of age-related changes in phenotypic variance affecting expression in humans and identified changes in variance with age as another process by which aging may be linked to disease.

Changes in variance with age occur either as a consequence of environmental exposures or as a result of changes in genetic regulation of gene expression. However, differences in expression levels within MZ twin-pairs must relate to the first explanation, as the individuals' genomes are identical. Therefore, and exploiting the twin design, we calculated the difference in expression between MZ co-twins (Additional Table S1). We have successfully used this strategy previously to classify genetic determinants of phenotypic variance in gene expression [15] and GxE interactions affecting allelic specific expression [16]. Here, we identified 42 genes where difference (discordance) in expression between MZ co-twins changed with age in at least one tissue (Figure 1C, Additional Figure S4 and Additional File 8). Of the 34 genes identified in skin, 11 also showed a decrease in variance with age (Additional Figure S5). This indicates that the observed change in variance for those genes was environmentally, and not genetically determined. However, for the remaining genes, either changing environments, concordant across MZ twins, or GxE interactions remain plausible explanations for the change in variance. In conclusion, changes in phenotypic variation with age can be attributed to different environmental exposures among the individuals and not only to a general decline in regulatory functions and increased genomic damage, as others have suggested [12].

### Age-related associations in expression are modulated by genetic variation

Changes in variance in expression with age could also be a result of gene-by-age interactions affecting expression (GxA), when the genetic regulation of expression changes with age [6, 24, 25]. To discover these effects, we searched for SNPs whose effect on expression levels depends on the age of the individual. It is well known that the power to discover such second order effects is much reduced compared to standard main effects; for this reason it is common to restrict the search space to those with known main effects, either genetic or on aging [26]. We concentrated on genes linked to age, as strong eQTL usually lie within promoter regions and as such are less likely to be environmentally influenced. We tested *cis*-GxA regulatory interaction effects with all SNPs in the cis window of 12,830 exons which were either 1) differentially expressed with age; 2) variance changes with age and 3) discordant in expression between MZ co-twins with age. After multiple testing corrections, we identified one significant GxA-eQTL, affecting the expression of *CD82* (Figure 2A). We also detected three GxA-eQTL among the genes that were discordant for expression in skin (Additional Figure S6 and Additional Files 9, 10, and 11). Despite the inherent challenges in identifying interaction effects, we here identify four GxA effects on gene expression with a relatively modest sample size. Given the many examples of GxA interactions reported in model organisms, we expected further studies with larger samples sizes to identify more examples.

**Figure 2A.**
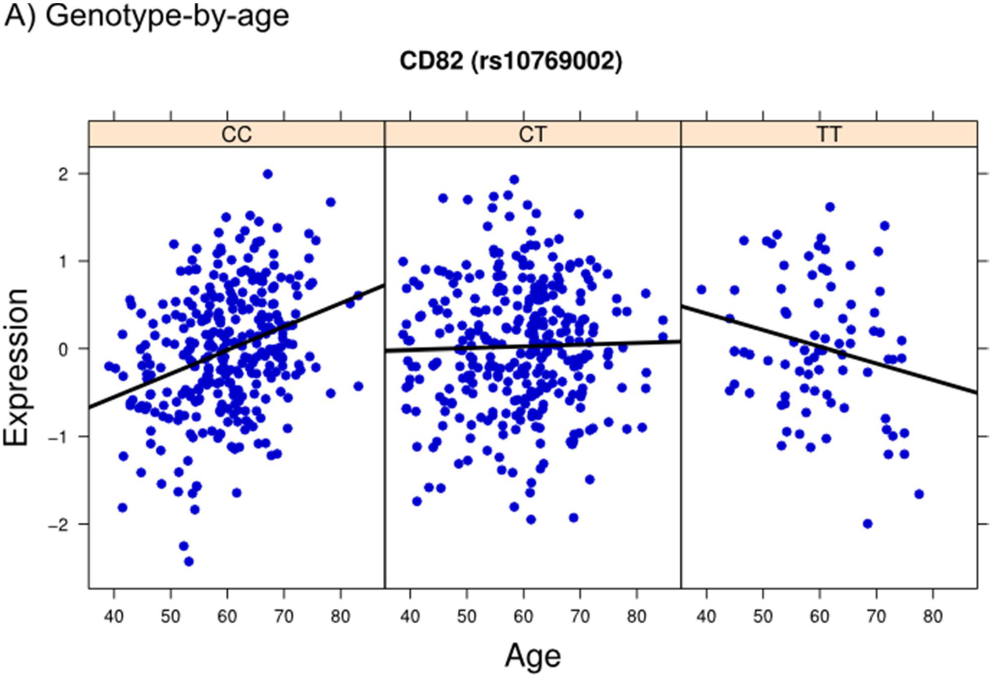
Interacting effects of aging on gene expression: The two plots show the effects of genotypes (eQTL) and methylation on gene expression can be modulated by age. A) The graph shows a genotype-by-age expression quantitative trait locus (gxa-eQTL) in fat tissue affecting the expression of the *CD82* gene. The expression of the reported exon increased with age in homozygous individuals for the CC alleles in rs10769002. Homozygous individuals for the alternative allele (TT) showed a decreased in expression with age.

**Figure 2B.**
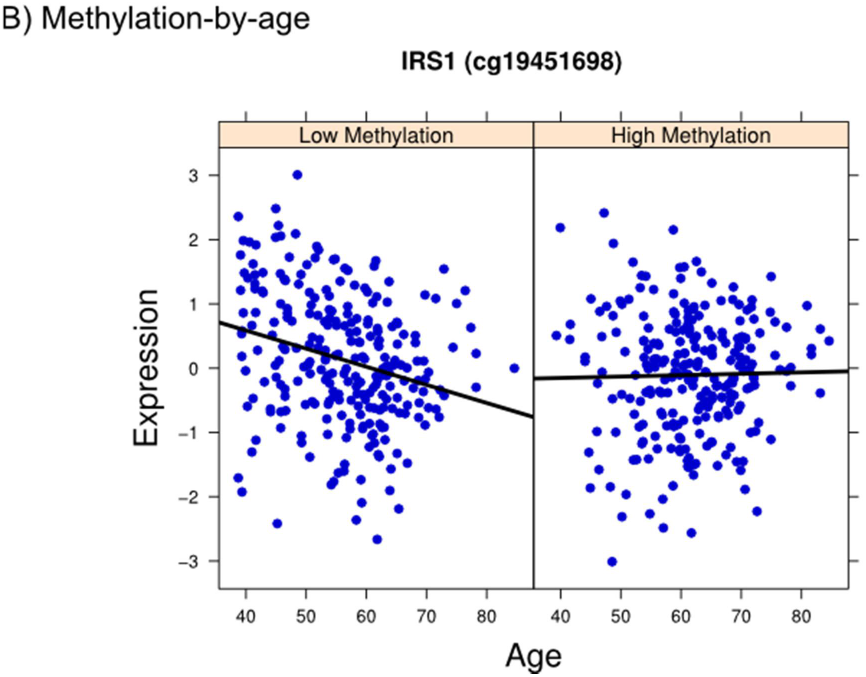
Interacting effects of aging on gene expression: The two plots show the effects of genotypes (eQTL) and methylation on gene expression can be modulated by age. B) The graph shows a methylation-by-age interaction affecting gene expression. The expression of the *IRS1* gene decreased with age in individuals with the methylation cg19451698 hypomethylated.

### The effect of age on methylation in fat tissue

Methylation levels and discordance in methylation between MZ twins has been shown to increase globally with age in promotor regions [27, 28]. Given our findings of genes with age-dependent changes in variance and discordance of expression, and the difficulties of identifying genetic effects responsible for those changes, we postulated that epigenetic drift could explain some of the age-related changes in expression. Using 552 Infinium HumanMethylation450 BeadChip methylation profiles generated from the same fat biopsies as the RNAseq data (Grundberg et al.), we identified 39,092 differentially methylated regions (DMRs) with age from the 370,731 array probes, 93.6% of which were hypermethylated with age (Figure 3, Additional File 12, methylation data was not available for other tissues). The proportion of DMRs was significantly larger than the proportion of age associated exons (10.54% compared to 1.4%, P value < 2.2e-16, X2 test). In total, 3,555 genes have an age-DMR near their TSS (<200 bp), of which 444 were also differentially expressed with age, suggesting methylation as a possible mechanism for age associated expression changes.

**Figure 3.**
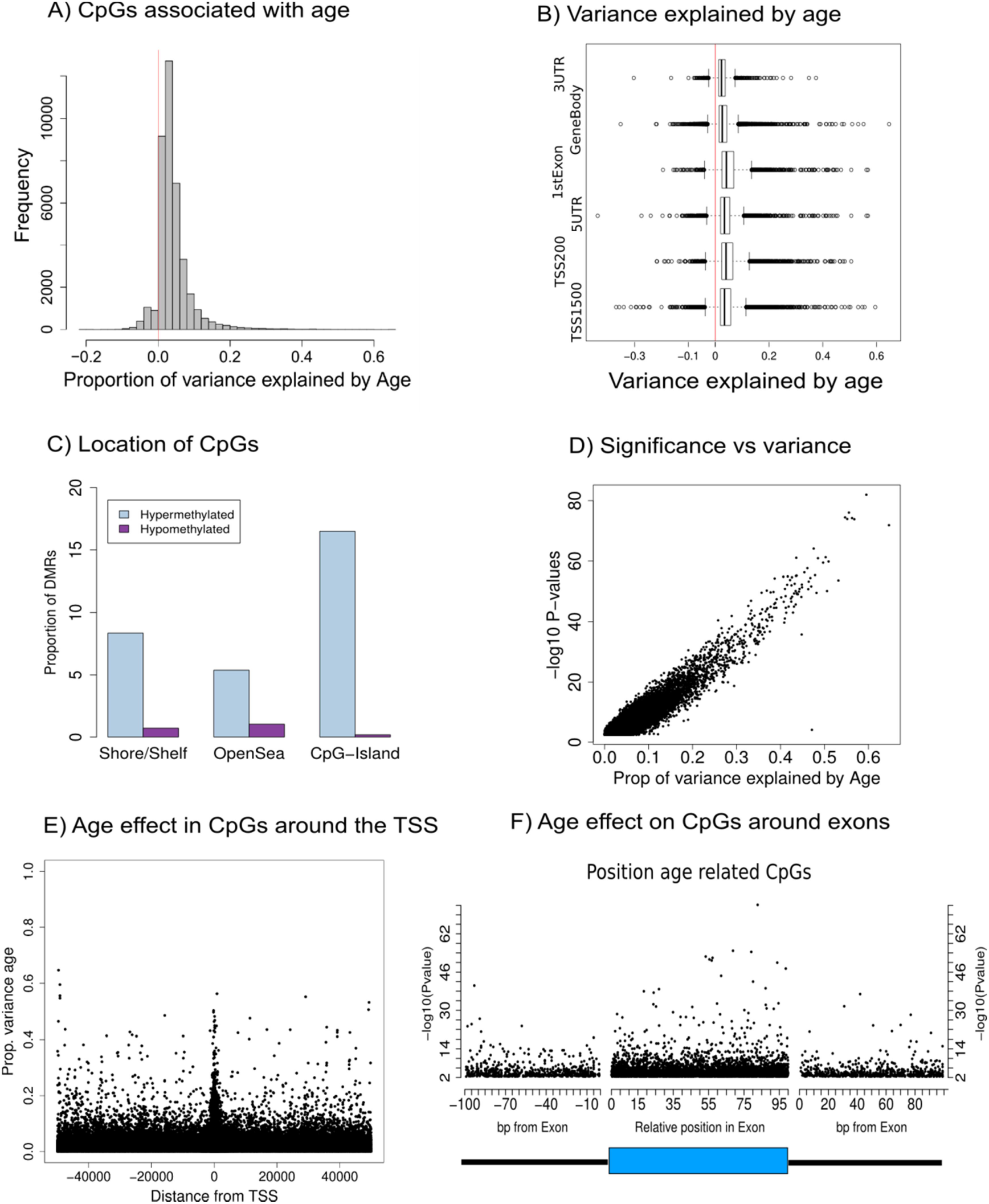
A) CpG islands showed mainly an increased in methylation with age, independent form the genomic position (B and C). D) The estimates for the proportion of variance attributed to age in methylation show that up to 60% of the variance in methylation would be attributed to age. E) The position of CpG markers respect to the near TSS from genes (only CpGs at <50Mb from the TSS are shown) show a larger effect of age on methylated regions near the TSS. F) The left and right panels show age-associated CpGs positions at the near 0-100bp at the 3’and 5’of each exon. The central panel show the relative position of the CpGs associated with age within each exon (blue box). The CpGs show higher associated with age in the exon 5’region, probably due to the proximity to the TSS of the genes.

In addition, we looked for associations between expression of exons with methylation probes within 200 bp of the TSS of the gene. Of 297,702 pairs of exon-methylation probes, we found 4,853 to be significantly correlated, 53% negatively and 46.91% positively correlated (Additional Figure S7 and Additional File 13). From those 4,853 exon-probe pairs, 16.8% of exons and 15.3% of methylation probes were differentially expressed or methylated with age. In conclusion, we observed a widespread and stronger effect of age on methylation than expression and a lower number of significant associations between expression and methylation.

To investigate whether interactions involving methylation markers are proxies for environmental factors explaining changes of variance with age in gene expression, we looked for interactions between methylation and age (methylation*age) affecting expression. Since only 3 genes had a significant association between variance in gene expression in fat and age at an adjusted P value < 0.05, we chose to relax our threshold to an FDR < 0.10 and test nine genes. We identified a Bonferonni significant methylation x age interaction effect on expression of *IRS1* at three methylation probes, most significantly *cg19451698* (*P* value = 6.6e-05, Figure 2B, Additional File 14). This significant interaction implies that the expression of the *IRS1* gene decreases with age in individuals where *cg19451698* is hypomethylated. Such an effect was not present in individuals with high levels of methylation in the same region. Homologs of *IRS1* and other members of the insulin/IGF-1 pathway are known to regulate longevity in model organisms, a function that may be conserved in humans due to their involvement in age-related diseases like type 2 diabetes. In summary, we detected an interaction between methylation and age affecting a gene expression showing how the effects of age can be modified by genetic and environmental factors.

### Age effects in expression are shared across tissues

Previous studies performed in multiple tissues identified a limited number of shared genes whose expression associated with age across tissues [3, 8, 29]. Similarly, of the 5,631 genes (36.67%) affected by age in at least one tissue, we were only able to identify five genes significantly associated to age in all the three primary tissues (Figure 4A, Additional Figure S8, Additional Table 5). We found 274 genes significantly associated with age in both fat and skin. This overlap was highly significant compared to what would be expected given independent gene sets in each tissue (*P* value < 1e-216, Fishers test), indicating the presence of a common signature of aging. However, it is important to notice that defining tissue-shared effects based on strict thresholds will underestimate the true sharing between tissues, particularly in blood which had reduced power to detect associations due to smaller sample size. Enrichment analysis, which can detect evidence of sharing which does not attain statistical significance by comparing the P value distributions across tissues, revealed shared age-related effects ranging from 21% to 60% (Figure 4C), with skin and blood showing the least overlap while fat and skin showed the most. This is considerably greater than the degree of enrichment observed in microarrays, 27%-28% between fat and skin [3]. We observed similar levels of tissue-shared age-related effects on splicing, with pairwise sharing ranging from 16% to 47%. In summary, our results indicate that global biomarkers of aging with effects across multiple tissues are prevalent.

**Figure 4.**
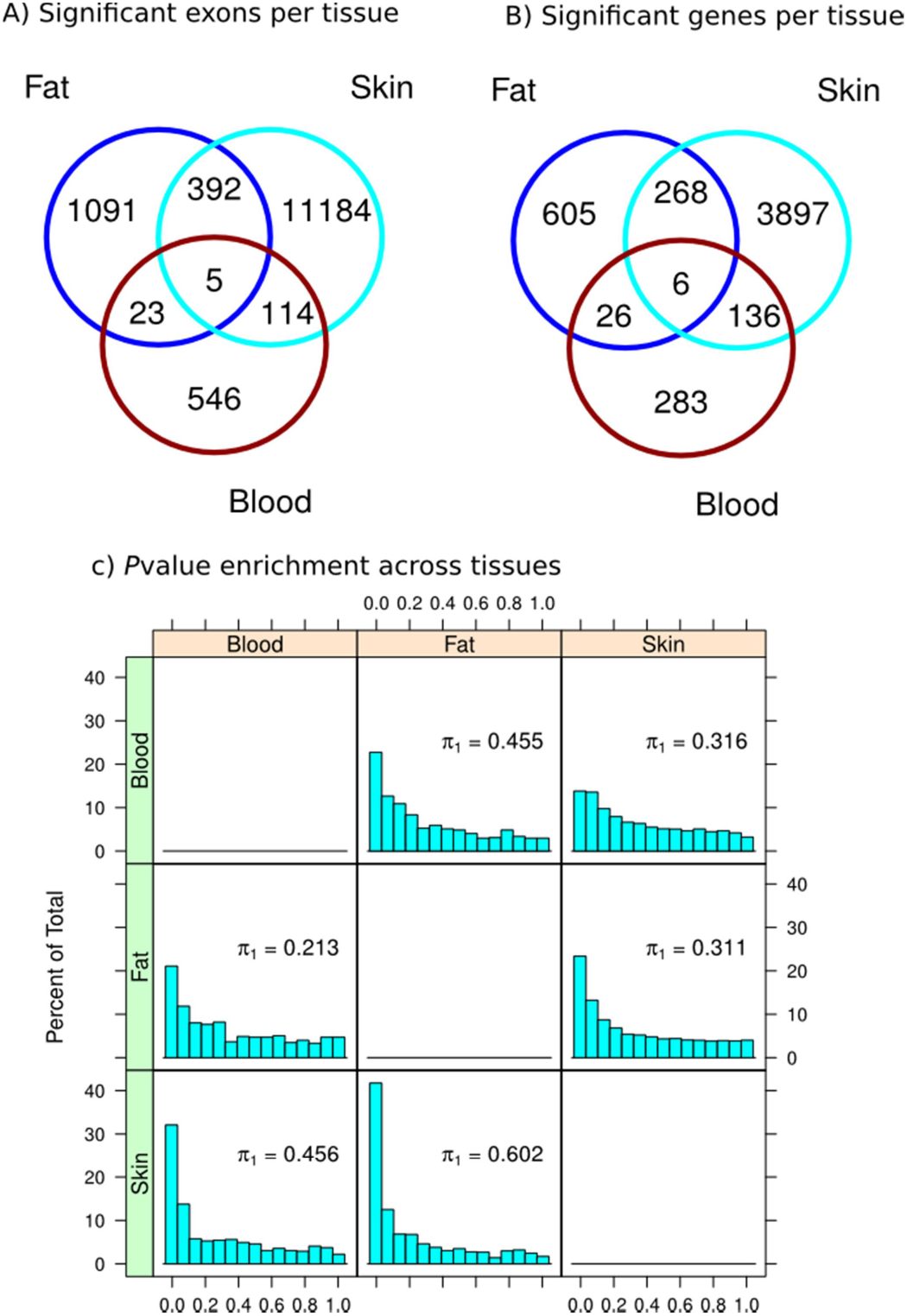
Tissue shared and specific effects of aging in gene expression changes with age. The top venn diagrams show A) the number of exons (left) and B) number of genes (right) significantly associated with chronological age in fat, skin and whole blood. Five exons were commonly associated to age in the three tissues. LCLs were not included, as only 7 exons were significantly associated with age. C) The *P* values of significant exons associated with age in one tissue were extracted from the analysis in the other tissues for enrichment analysis (π1). The histograms show the *P* values for association between expression and age in one tissue (left, green color) if the exons were significantly associated exons in another tissue (top, orange color). As show in the graphs, age-related signals detected in fat shared an estimated 60.2% of the age effect signal skin tissue and 45.6% with blood.

## Discussion

The association between aging and disease has been extensively demonstrated by epidemiological and GWAS studies [2, 30], but the association between specific genes with a disease in the context of the aging process remains elusive. We here identify chronological age-related changes in the expression of thousands of genes. Gene expression is an important endophenotype that underlies susceptibility to many common diseases and could mediate the effect of age on disease. To illustrate the many different effects of age in the expression here investigated we present here two examples of disease-related genes: *APOE* and *LMNA*. *APOE* has been associated with Alzheimer and cardiovascular diseases, and genetic variants within the *TOMM40/APOE/APOC1* locus have been linked to longevity [31]. Our analysis showed that the expression of multiple exons and links of *APOE* change with age in skin tissue, producing different isoforms that can potentially induce changes in the activity of the gene (Figure 5). Based on current models for *APOE* transcripts, we expect all transcripts to include exons 2, 3 and 4, with the possibility of including exon 1 given the use of an alternative TSS. However, the significant increase in the number of reads linking exons 2 and 4, and reduction of those linking exons 3 and 4, indicates that transcripts skipping exon 3 are more common in older individuals. Such a transcript has not being reported as functional, for which we hypothesize that the *APOE* gene produces aberrant and likely non-functional transcripts in older individuals after exon skipping events. Whether this reduces the overall expression of functional transcripts, leading to lower levels of circulating apoliprotein E, requires further testing. Additionally, we previously reported an eQTL affecting the expression of *APOE* in skin and fat tissues [16]. The lead SNP of this eQTL (*rs439401*) has been implicated in triglycerides metabolism, Alzheimer's and cardiovascular diseases [32]. We here find a nominally significant gene x age interaction eQTL for this SNP (*P* = 0.014), suggesting the regulatory effect of this SNP on *APOE* expression is dependent on age. Given the strong association between *APOE* and age-associated diseases, such an effect could potentially be mediated by age-dependent changes in genetic regulation of *APOE* expression.

**Figure 5.**
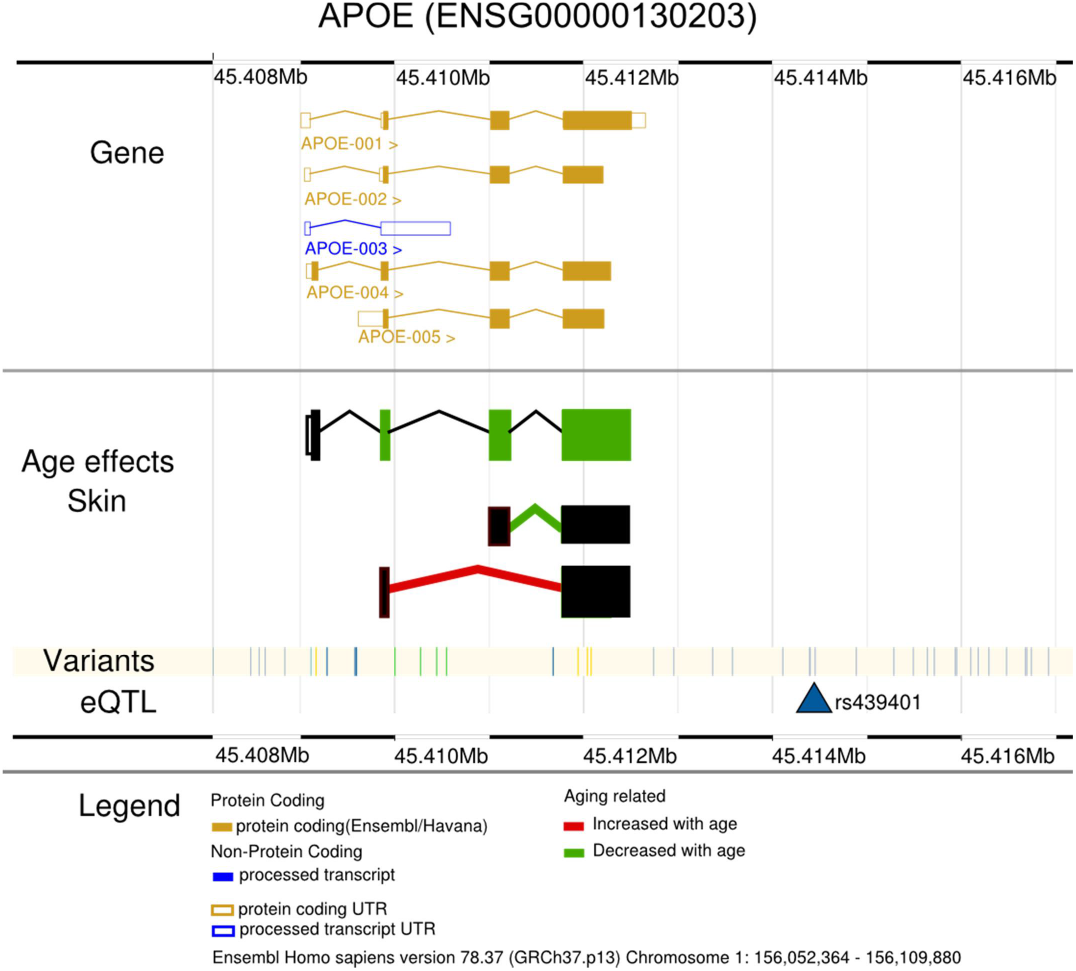
Structure of the APOE gene. Mutations in this gene have been associated with alterations in fatty acid metabolism and cardiovascular diseases. Polymorphisms in and near the gene has been associated with Alzheimer and cardiovascular diseases. The gene produces multiple protein coding transcripts variants (yellow) and non-coding processed transcripts (blue). In the skin tissue, three exons and one link decreased their expression with age (green coloured exon and link between exons); and one link increase its expression (red coloured link). Furthermore, we detected one eQTL (rs439401) affecting the expression of the gene.

The second example we choose to highlight here involves the *LMNA* gene (Figure 6), which is causal of the Hutchinson-Gilford progeria syndrome. This syndrome is characterized by accelerated aging features as a consequence of the accumulation of a truncated progerin isoform of *LMNA.* The quantity of the progerin transcript increases with age in normal cells [33], with its protein known to accumulate in human skin in an age-dependent manner [34]. We identified age-related changes in expression of *LMNA* exons (FDR < 0.1) and links (FDR < 0.05) consistent with the production of different alternative isoforms in an age-dependent manner. With increasing age, we observe an increase in the number of reads corresponding to exon 7 in the plot (red reads in Figure 6), and a decrease in the number of reads corresponding to exon 3. This by itself suggests that transcripts which include exon 7 but skip exon 3 are more prevalent in older individuals. This is consistent with our exon-link data, in which reads linking exon 2 to exon 4 (thereby skipping exon 3) were also more abundant in older individuals. Furthermore, we found an eQTL affecting *LMNA* expression in skin, blood and LCLs tissues, the peak LCL eQTL (*rs915179*) has been previously linked to exceptional longevity in humans [35, 36].

**Figure 6.**
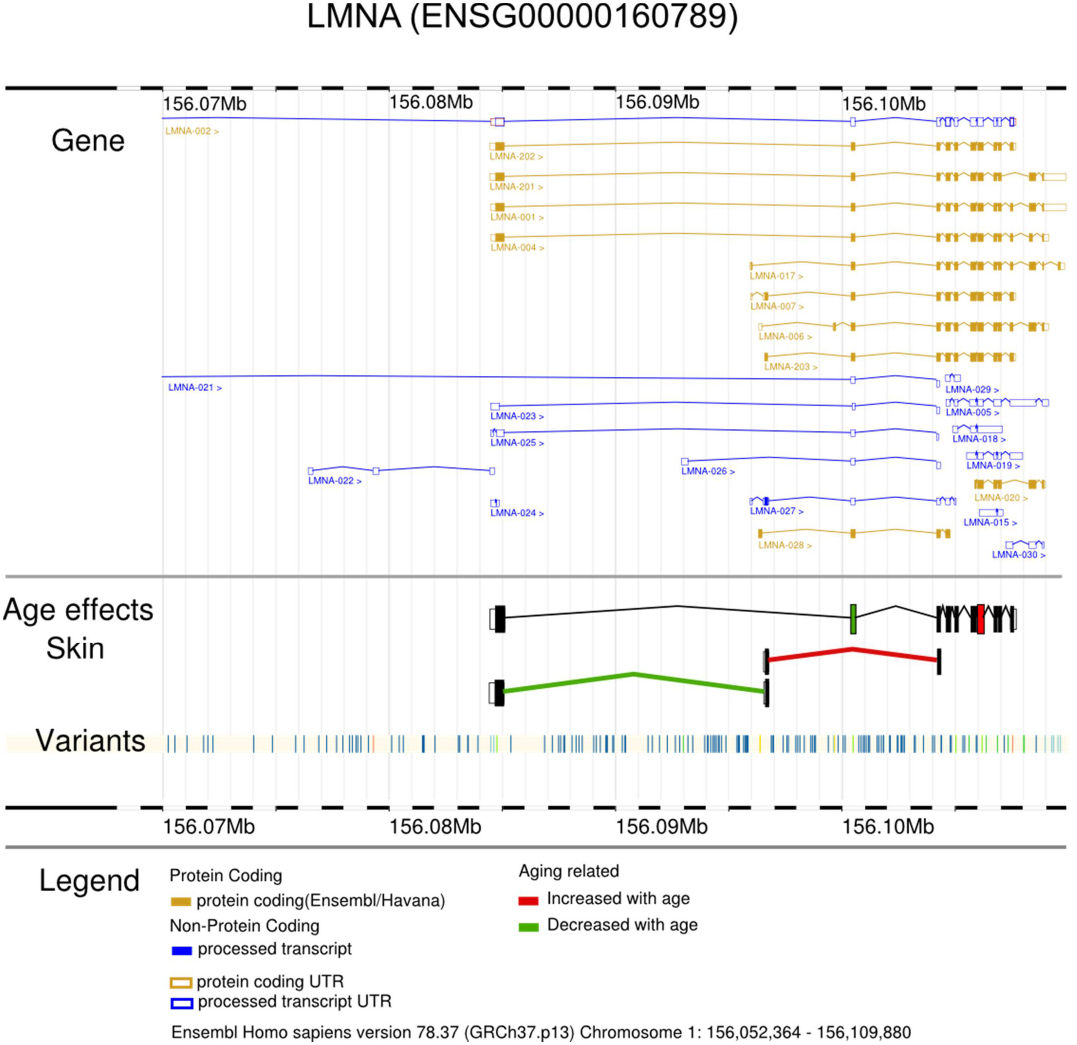
Structure of the LMNA gene. Mutations in this gene has been associated with multiple diseases, including the Hutchinson-Gilford progeria syndrome, characterize by accelerated aging features. The gene produces multiple protein coding transcripts (yellow) and non-coding processed transcripts (blue). In the skin tissue, two exons were affected in their expression by age by increasing expression (red coloured exon, corrected Pval < 0.1) and decrease expression with age (green coloured exon). Furthermore, two links were significantly associated with age in their expression (corrected Pval < 0.05). Our results suggested an increase in the production of isoforms using alternative 5’.

Both examples presented for *APOE* and *LMNA* show age related changes in splicing of expression in genes linked to age-dependent diseases, as well as demonstrating how genetic variation can ameliorate these changes. However, it is important to take into account that the gene models included on our figures and used as references are based on GENCODE annotated transcripts. Some of the annotated transcripts may not necessarily translate into proteins and the used of annotated gene models means we do not account for the reported presence of aberrant or incomplete transcripts [37] We also note that some of the effects of age on gene expression, splicing and methylation will be mediated by age-related changes in cellular composition. Future studies should investigate the extent to which this is the mechanism underlying our observed age-related changes in expression.

An under-studied hypothesis liked to age-related changes propose that an increase in age-related disease prevalence is a consequence of a loss in regulatory capacities in aging organisms, which manifested in an increase in phenotypic variance with age [10, 11]. This theory assumes that only an increase in variance can be associated with disease. However, a decrease in expression variance could indicate a reduction in the ability to respond to the organism's environment or to reduced regulation of expression, either of which could lead to disease development. In contrast with reported patterns of increased expression variance with age, we mainly identified individual genes with an age-related decreased variance. This contradicts the expectation of a stochastic increase of the phenotypic variance with age due to reduced regulatory capabilities. Three factors may induce changes in phenotypic mean or variance: genetic variation, environmental variation or an interaction between the two. Genes and pathways associated with longevity and age-related changes are often strongly regulated in older organisms with low levels of stochasticity and higher levels of heritability [6, 11, 38]. In this study, the exons associated to age were highly heritable (Additional Figure S9, Additional Table S6 and Additional file 3) suggesting, as previously reported, that age modulates genetic regulation of expression [6].

We attempted to identify genetic and environmental factors involved in the changes of variance with age by testing for GxE interactions. We were able to identify a significant gxa-eQTL in fat tissue acting on the gene *CD82* (*rs10769002*). This gene is associated with tumor progression as it codes for a metastasis suppressor glycoprotein highly correlated with *p53* and which increase in expression have been associated with overall better survival to cancer [39]. In our analysis we observed that individuals homozygous for the reference allele increased gene expression with age compared to the alternative allele. Therefore, it is possible that the alternate allele in *rs10769002* may be a risk factor for some types of cancer in older individuals. Three other examples were identified in skin tissue for genes also previously implicated in cancer and metabolism. In conclusion, we identify changes in phenotypic variance with age that would be explained by GxE and changes in regulation, suggesting that damage accumulation is not the only explanation to the observed change in phenotypic variance with age in many other phenotypes. Moreover, we show that the study of phenotypic variance with age in gene expression may identify new candidate genes relevant for age-related diseases.

For a more mechanistic understanding of how age may affects expression, we investigated and detected a widespread effect of age on methylation. The effect of methylation was stronger than the observed effect of age on gene expression, although both age effects were small compared to the relative influence of genetics. Our search for interactions that would explain changes in variance with age identified *IRS1* as a gene whose expression changes as a consequence of an age-methylation interaction. The *IRS1* gene has been associated to type 2 diabetes, an age-related disease and it has also been found to have diabetes associated DMRs nearby [40]. Our results suggest that although methylation changes are strong markers for the aging process, their influence on expression changes with age may be only relevant for a small percentage of genes. We also observed that most age-DMRs were hypermethylated in our study, while there is currently no consensus in the literature on whether age-DMRs are primarily hypermethylated or hypomethylated in a particular tissue [41]. In conclusion, our search to explain results on global changes in phenotypic variance with age indicates that increase or decrease in expression regulation jointly with the accumulation of environmental exposures may often be observed as multiple GxE interactions.

## Conclusions

In summary, we have performed a large human transcriptomic study of aging, in a multi-tissue dataset. We took advantage of studying aging in a healthy cohort to identify genes with expression associated with age. Since age-related changes are highly unlikely to be confounded with disease prevalence in this dataset, genes previously linked to disease and here associated with age are more likely to be causative of disease. Therefore, the thousands of age-related genes identified in this paper are of high interest to researchers studying these diseases. In addition, we found that the shared effect of aging in humans across four tissues as well as the number of affected genes is larger than previously reported. We also quantified the global effect of age in gene expression reporting a small influence in expression (median variance explained by age is between 2.2% and 5.74%). When compared to the large global effect of genetic factors on gene expression, the low age-related values may explain the difficulties in identifying biomarkers of aging in gene expression, and highlight the value to account for genetic variation when considering individual level biomarkers. On the other hand, we observed a larger global effect of age in methylation levels in the same fat samples, with age explaining up to 60% of the variance observed in methylation levels in some regions. However, we observed a low number of associations between expression and methylation, suggesting that the relationship between both phenotypes and age-related changes may be independent for most genes. Moreover, we have shown that age alters gene expression in multiple complex ways, including total expression, variance, splicing and genetic regulation. Many of the genes subjected to age-related changes in expression have been linked to age-related diseases, highlighting the need for future studies into the relationship between age-related changes in gene expression and its regulation, and age-related diseases. This is particularly relevant for genome wide association studies (GWAS) where eQTLs are routinely used to identify target genes of genetic variants without accounting for the effects of age.

## Methods

### Study design

The sample collection, and mRNA extraction has been described in detail in [42]. In sort, 856 Caucasian female individuals (336 MZ and 520 DZ twins) from the TwinsUK Adult twin registry [43] were recruited with a ranged age from 39 to 85 years (mean 59 years). Samples were prepared for sequencing and processed as described in [15] and [16]. The number of monozygotic (MZ), dizygotic (DZ) and unrelated individuals (individuals with no relatives in the dataset) included in the final analysis per tissue are described on Additional Table S1

### Exons and links quantification

The 49-bp sequenced paired-end reads were mapped to the GRCh37 reference genome (The International Human Genome Sequencing Consortium, [44]) with BWA v0.5.9 [45]. We use genes defined as protein coding in the GENCODE v10 annotation [46], removing genes with more than 10% zero read count in each tissue. For the analysis presented in this paper, only exons from protein coding genes and LincRNAs from verified loci (level 1) and manually annotated (level 2) were investigated. We calculated the relative quantification of splicing events using Altrans [22]. Sequencing of each sample produced different number of reads, therefore read counts assigned to links and exons needed to be scaled to 10 million reads to allow comparisons across samples.

Additional Table S3 show the total number of exons and genes sequenced per tissue, as well as the total number of exons, genes used in the analysis here presented. We quantified expression at the level of exons as exon-level quantifications provide both a greater resolution than gene-level quantifications and are more biologically relevant. Gene-level analysis assumes that effects operate on the mean level of all transcripts, whereas exon-level analysis allows the identification of effects due to changes in mean expression of subsets of transcripts or changes in alternative splicing.

### Genotying and imputation

Genotyping of the TwinsUK dataset (N = ∼6,000) was done with a combination of Illumina arrays as described in [15, 16, 42]. Samples were imputed into the 1000 Genomes Phase 1 reference panel (data freeze, 10/11/2010) [47] using IMPUTE2 [48] and filtered (MAF<0.01, IMPUTE info value < 0.8).

### Splicing junction quantifications

We calculated the relative quantification of splicing events using Altrans [49]. The method makes use of mate pairs mapped to different exons to count “links” between two exons based on the GENCODE v10 annotation for level 1 and 2 from protein coding genes and lincRNA. Exons that overlap were grouped into “exon groups” to identify unique portions of each exon from an exon group. The unique portions were used to assign reads to an exon. The quantitative metric produced by Altrans is the fraction of one link's coverage over the sum of overages of all the links that the primary exon produced. The values range from 0 to 1, representing the proportion of a give link among all the links produced by the primary exon. The metric is calculated in 5′-to-3′ (forward) and 3′-to-5′ (reverse) directions to capture splice acceptor and donor effects respectively. Additional Table S4 show the total number of links identify per tissue, as well as the total number of links per gene detected.

### Age effects on mean exon expression and links

Rank normalized reads per exon or links were used to assess the age effect on exon expression mean. A linear mixed model was fitted to examine age effect on gene expression in R [50] with the lmer function in the lme4 package [51]. Confounding factors in all models included fixed (primer insert size, GC content mean and batch (only for blood samples)) and random effects (primer index, date of sequencing, family relationship and zygosity). The *P* values to asses significance for age effect were calculated from the Chi-square distribution with 1 degree of freedom using likelihood ratio as the test statistic. A set of 100 permutations were used to adjust for multiple testing. Expression values were permuted while maintaining samples from twin pairs together.

To account for the fact that genes have different numbers of exons, we stratified the genes into 16 groups based on number of exons and applied the multiple testing correction separately to each group. In this way we control the false discovery rate while simultaneously ensuring that we do not see a higher proportion of false positive results for genes with more exons. The adjusted P values, controlling the FDR, were calculated as the proportion of permuted statistics more significant than the P value calculated on real data, divided by 100 (for the number of permutations). P values < 0.05 were considered significant. A gene was considered as significantly affected by age if the expression of at least one exon was significantly associated to age.

### Tissue shared effects

For each pair of tissues comparison we extracted P values of exons in one tissue (e.g. skin) from significantly age associated exons in other tissue (e.g. fat). The P values distributions were used to assess the enrichment of age associated exons in other tissues. Analysis were performed in largeQvalue [52], an implementation of the R statistical software qvalue package [53], for large datasets.

### Number of genes expressed with age

Raw FPKM read counts were used to identify the number of genes expressed per individuals. A gene was considered expressed with FPKM read counts > 0.2. The numbers of expressed genes were rank normalized and used to assess the age effect on number of genes expressed. A linear mixed model with number of genes expressed per samples as response variable was used to assess the association between number of exons expressed and age. Confounding factors in all models included the same fixed and random effects as used before. *P* values were calculated from the Chi-square distribution as before.

### Age effect on variance of gene expression

Residuals removing from technical covariates and family structure were used to assess the association for variance and age per tissue. Residuals were extracted from a linear mixed model fitted with the lmer function in the lme4 package [51] using R. Confounding factors in all models included fixed and random effects as detailed above. The residuals were fit on a loess function including age as response variable. Residuals from the loess regression were squared root to give a measure of the distance from the mean expression with age. A Spearman correlation test between this 'distance' and the age was used to asses evidences for an age effect on variance. Multiple testing corrections were performed as described for the expression association with age with 100 permutations.

### Age effect on discordance of gene expression

Residuals from expression after removing only technical covariates were used to assess the change in discordance of gene expression with age per tissue from complete MZ pairs of twins (Additional Table S1). Association with age was assessed by regressing the maximum expression of each twin pair on the expression of the sibling plus age to detect whether the relationship between maximum and minimum expression was conditional on age. Multiple testing was assess using 100 permutations and as described for the expression association.

### Fat methylation analysis

Methylation data were downloaded from ArrayExpress, accession number E-MTAB-1866. The authors used Infinium HumanMethylation450 BeadChip (Illumina Inc, San Diego, CA) to measure DNA methylation in 648 female twins. Further details of experimental approaches can be found here [54]. Since that publication we adopted a different QC procedure (BMIQ) [55] that identifies low quality samples and corrects the technical issues caused by the two Illumina probe types. Therefore, the number of samples after the new quality assessment was 552, of which 516 also had fat expression measured with RNA-seq. The methylation data was also background corrected. DNA methylation probes that mapped incorrectly or to multiple locations in the reference sequence were removed. Probes with >1% subjects with detection P-value > 0.05 were also removed. Subjects with more than 5% missing probes were also removed. All probes with non-missing values were included.

Differential methylation with age was investigated for probes around the 50,000 bp from the TSS of genes included in the age analysis, which give us a total of 370,731 probes tested from a total of 541,369 CpGs probes on the 450K array. A linear mixed model was fitted to examine age effect on gene expression as in previous analysis. Confounding factors in the models included fixed (beadchip, BS conversion efficiency and BS-treated DNA input) and random effects (family relationship and zygosity). Multiple testing was assessed using 100 permutations. Methylation expression association was tested using expression residuals after removing technical covariates and family structure using a linear model in R with. 100 permutations were used to correct for multiple testing.

### Effect sizes and heritability analysis

We calculated effect size of age in expression and methylation from the normalized data and as a proportion of variance attributed to age over the total variance in exon expression. We also calculated the variance attributed to additive genetic effects, common environment and unique environment. Variance components were calculated from a linear mixed model, as previously described in [42], and [56] using all available complete twin pairs per tissue (Additional Table S1). The model was fitted as described above.

### Genotype-by-age and methylation-by-age interactions

Expression residuals removing from technical covariates and family structure were used to assess the association of exons and genetics variance interacting with age. To identify genotype-by-age interactions affecting gene expression we performed a linear regression of the residuals of each exon on the SNPs in a 1Mb window around the transcription start site for each gene, using a linear model in R. Only SNPs with MAF >= 0.05 were tested. We used 10 permutations to assess the significance of the interactions for exons with age-related effects, namely mean expression changes, variance changes and discordant effects. We used a similar strategy as used by [57] and based on [58]. A linear model with main effects but without an interaction term was used to extract residuals for each exon-SNP association test. The residuals were permuted (10 times) and used in a linear association with a model for the interacting term (gxa). *P* values from this analysis were stored and used to adjusted *P* values correcting for the number of exons per genes, as described before.

Methylation-by-age interaction analysis used expression and methylation residuals after removal of technical covariates and accounting for family structure. A linear model was used to test the association between expression and methylation levels with age. Significant associations were considered those with a P value < 1.0e-4 (Bonferroni correction).

### Code

Additional File 15 contains code use for the analysis presented in this manuscript.

## Declarations

### Ethics approval and consent to participate

This project was approved by the ethics committee at St Thomas' Hospital London, where all the biopsies were carried out. Volunteers gave informed consent and signed an approved consent form prior to the biopsy procedure. Volunteers were supplied with an appropriate detailed information sheet regarding the research project and biopsy procedure by post prior to attending for the biopsy.

### Data availability

RNAseq data is available in the European Genome-phenome Archive (EGA) under the accession EGAS00001000805. Methylation data is available in ArrayExpress, accession number E-MTAB-1866.

### Contributions

AV, AAB, AB and MND analyzed genotype and expression data as well as developed the methodology. AV, PCT and JTB analyzed methylation data. AV drafted the manuscript which contributions from all authors. AV, ETD, TDS and KSS designed the study. All authors read and approved the final manuscript.

## Acknowledgements

Some computations were performed at the Vital-IT (http://www.vital-it.ch) Center for high-performance computing of the SIB Swiss Institute of Bioinformatics.

## List of Additional files

Additional file 1: Summary statistics for exons association with age. Each sheet contains the pvalues for one tissue.

Additional files 2: Output from the functional analysis of differentially expressed genes with age using David. Only significant results (FDR < 0.05) are included. Each sheet contains the pvalues for one tissue.

Additional files 3: Results for variance decomposition per exon (variance explained by age and heritability), in fat, skin, whole blood and LCLs. Each sheet contains the pvalues for one tissue.

Additional files 4 and 5: Summary statistics for links (reads between exons) association with age. Tab delimited files, one per tissue and direction (reverse, forward): fat and skin. Each sheet contains the pvalues for one direcction.

Additional files 6: Output from the functional analysis of differentially expressed links with age using David. Only significant results (FDR < 0.05) are included. Each sheet contains the pvalues for one tissue (skin and fat).

Additional files 7: Summary statistics for variance of expression association with age. Each sheet contains the pvalues for one tissue: fat, skin, whole blood and LCLs.

Additional files 8: Summary statistics for discordance of expression between MZ twins and its association with age. Each sheet contains the pvalues for one tissue: fat, skin, whole blood and LCLs.

Additional files 9: Summary statistic for gxa-eQTL in differentially expressed exons in fat and skin. Only the best SNP-exon association per exon in included.

Additional files 10: Summary statistic for gxa-eQTL in exons with a significant change in variance with age in fat and skin. Only the best SNP-exon association per exon in included.

Additional files 11: Summary statistic for gxa-eQTL in exons discordant for expression between MZ twins in fat and skin. Only the best SNP-exon association per exon in included. Additional files 12: Summary statistics for methylation association with age in fat tissue.

Additional files 13: Summary statistics for methylation association with expression. Additional files 14: Summary statistics for methylation interaction with age.

Additional files 15: R code to perform the analysis discussed in the paper.

---

**Additional Figure S1.**
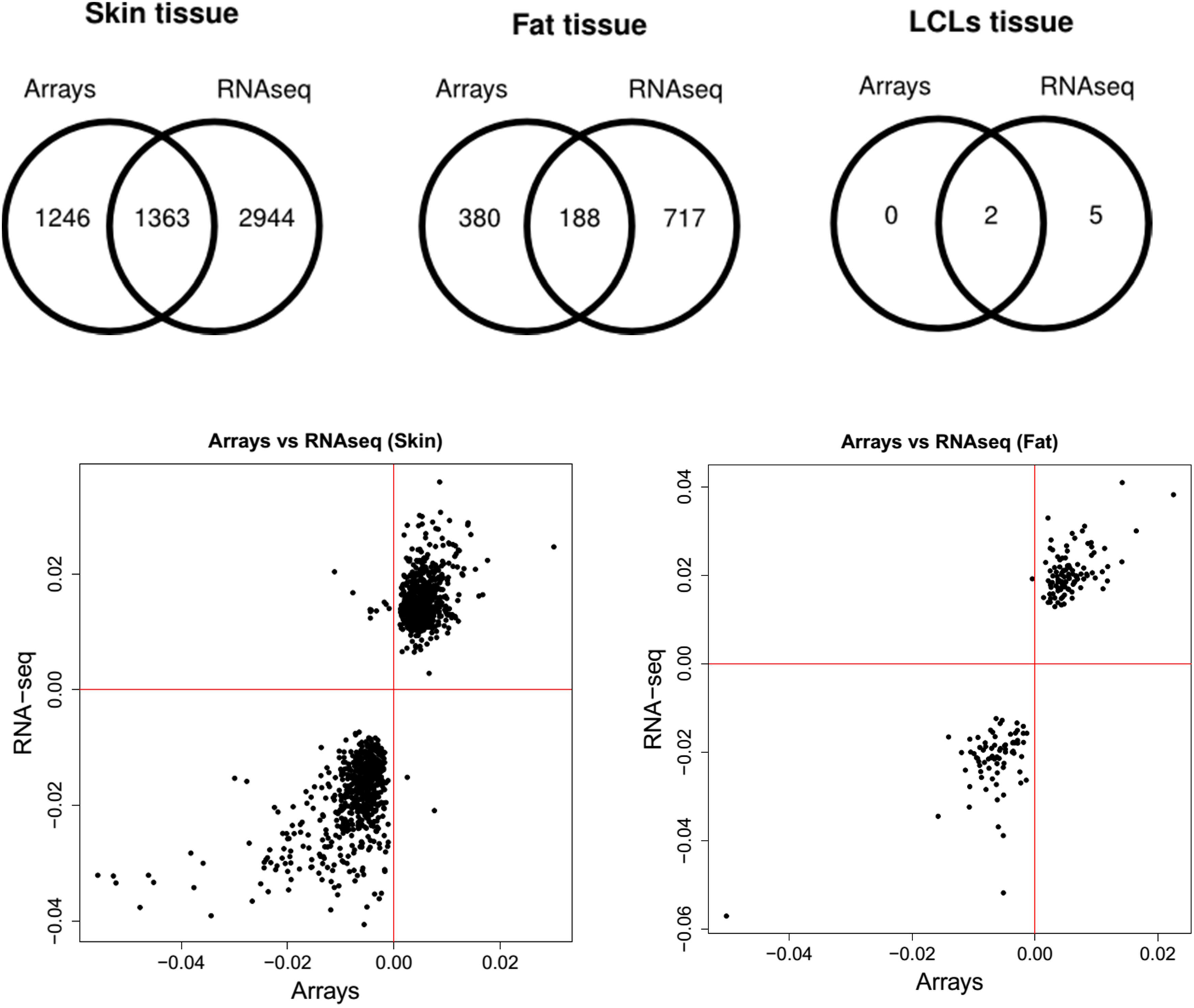
Top: Venn Diagrams per tissue comparing differentially expressed genes in array expriments (Glass et al, 2012) and gene swith at least an exon differentially expressed with age from RNA-seq data. Bottom: we compared the direction of effect of significantly affected genes with age in both technologies in skin tissue and fat tissues. The plot shows how RNAseq data replicates the same direction of effect for age in the majority of the significant genes in both technologies, which should be expected as they relate to quantifications of the same underlying phenotype in the same samples. For the very few genes with opposite effects, alternative splicing may explain the differences.

**Additional Figure S2.**
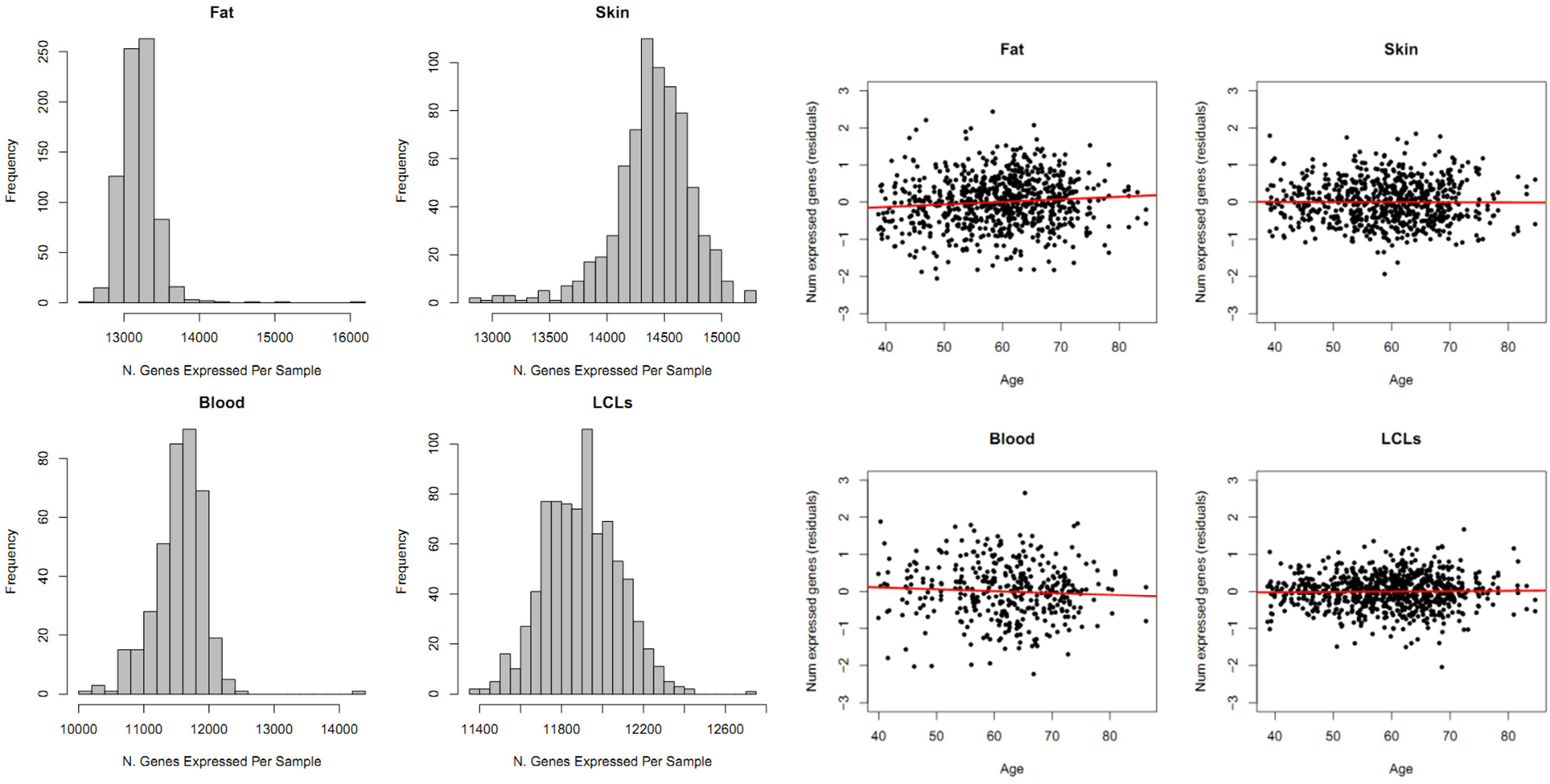
On the left, histograms with the frequency of number of genes expressed per sample in each tissue. On the right, we show the linear association between the number of expressed genes in each tissue (FPKM > 0.2) and age. Only the number of genes expressed was significantly associated with age in fat tissue (*P* value = 0.0026).

**Additional Figure S3.**
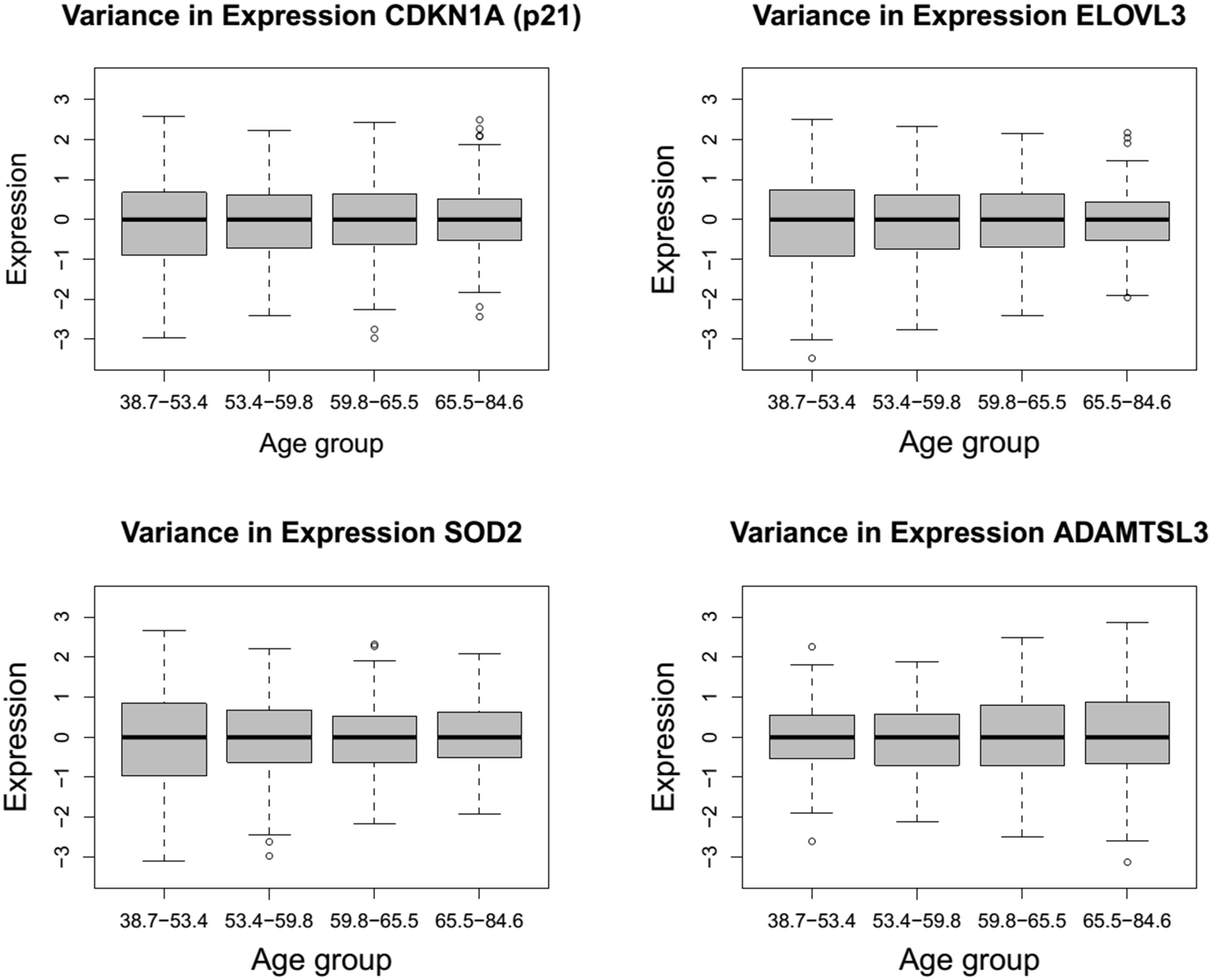
Examples for significant effect of age in variance of exon expression in skin. For this plot, individuals were grouped by ages, as indicated on the x-axis, with their expression values for the genes centered by the median expression, showing a decrease (top and bottom left plots) and an increased (bottom right) in variance with age.

**Additional Figure S4.**
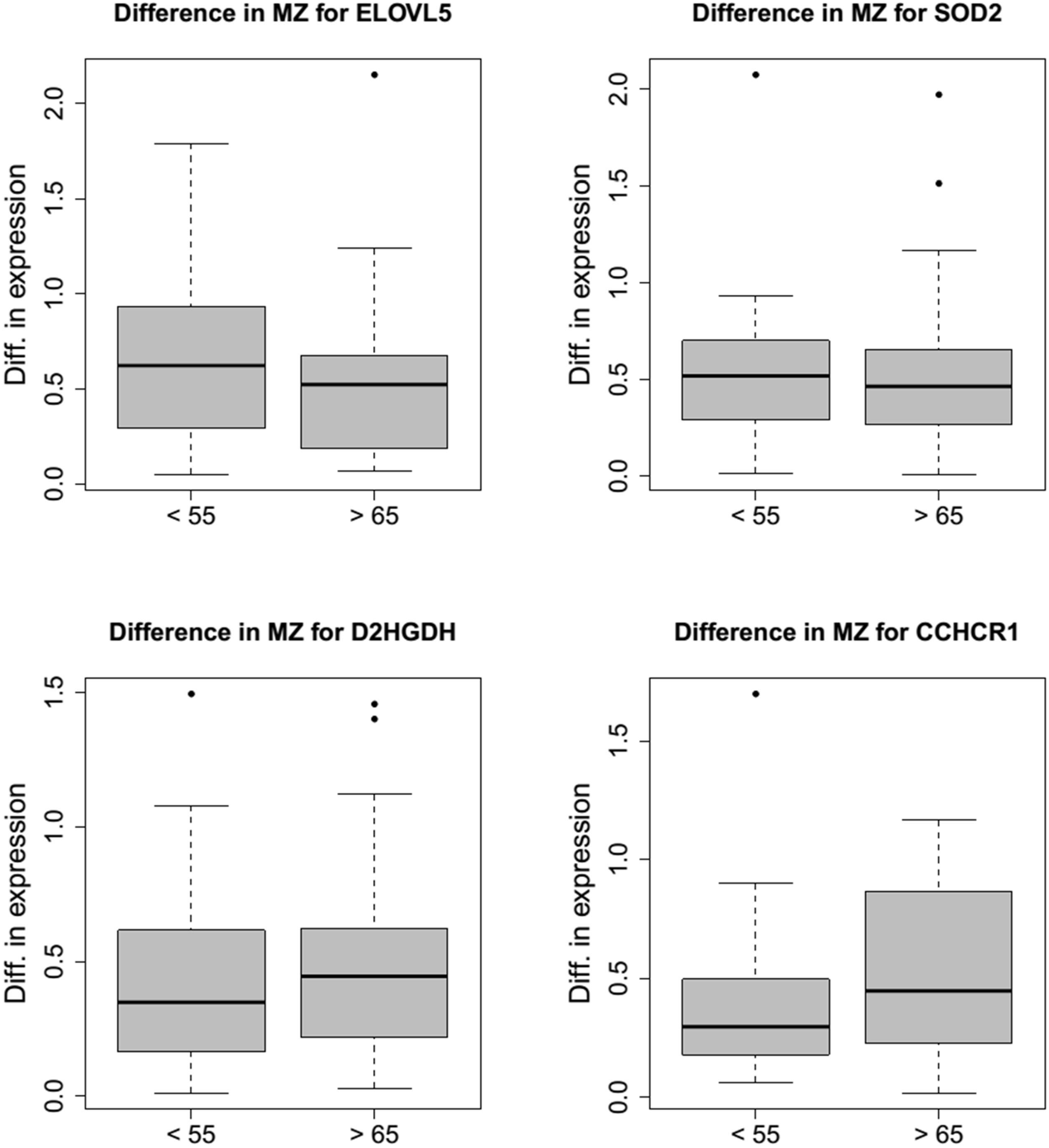
Barplots showing the difference in expression between MZ twins younger than 55 years old and older than 65 years old for genes with a significant decrease (top plots) and increase (bottom plots) in variance.

**Additional Figure S5.**
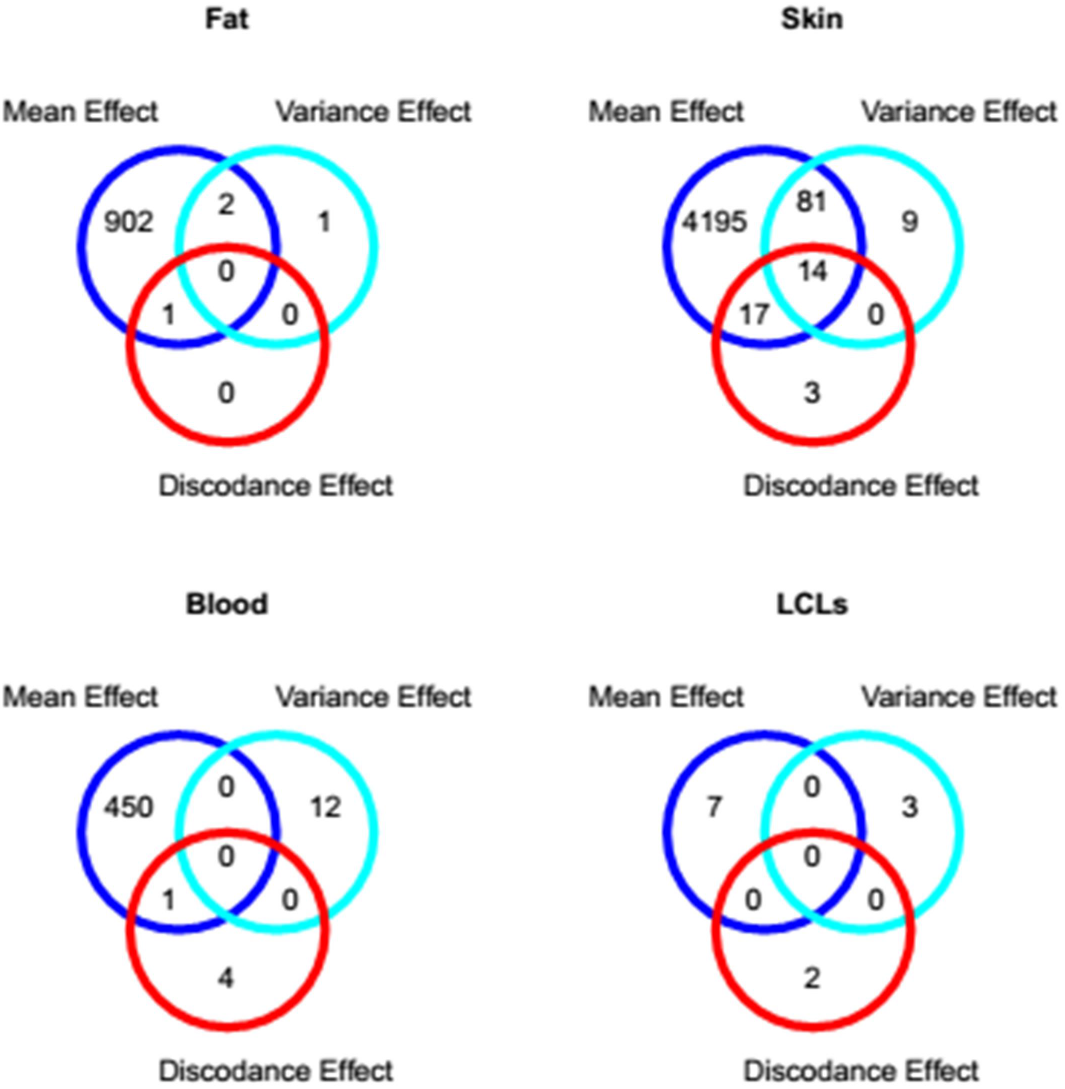
Venn diagram showing the overlap in genes with at least one exon affected by changes in the mean (DE genes), variance and differences between MZ twins (discordance) in each tissue.

**Additional Figure S6.**
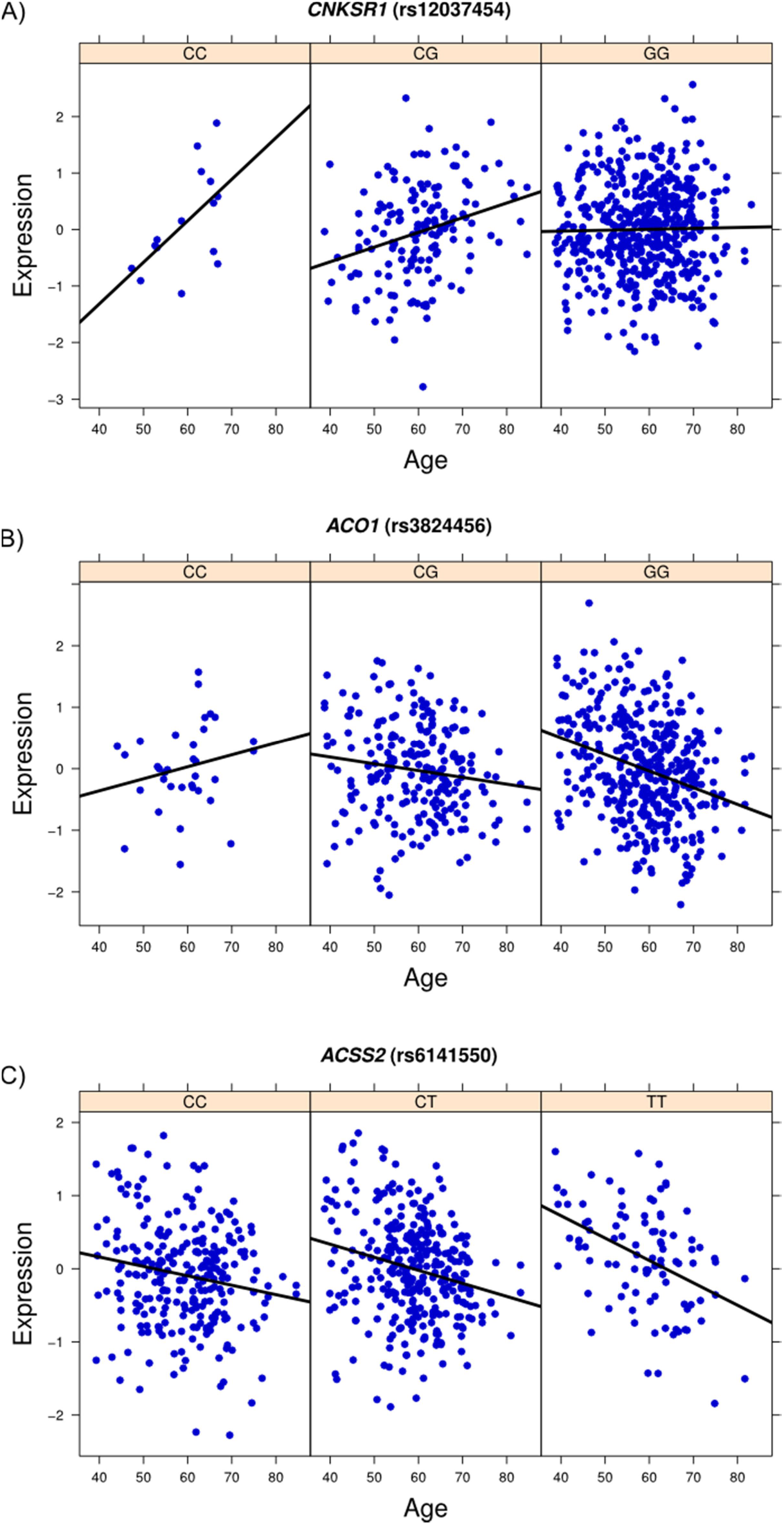
**Interacting effects of aging on gene expression:** All the graphs show a genotype-by-age expression quantitative trait locus (gxa-eQTL) in skin tissue affecting the expression of three genes.

**Additional Figure S7.**
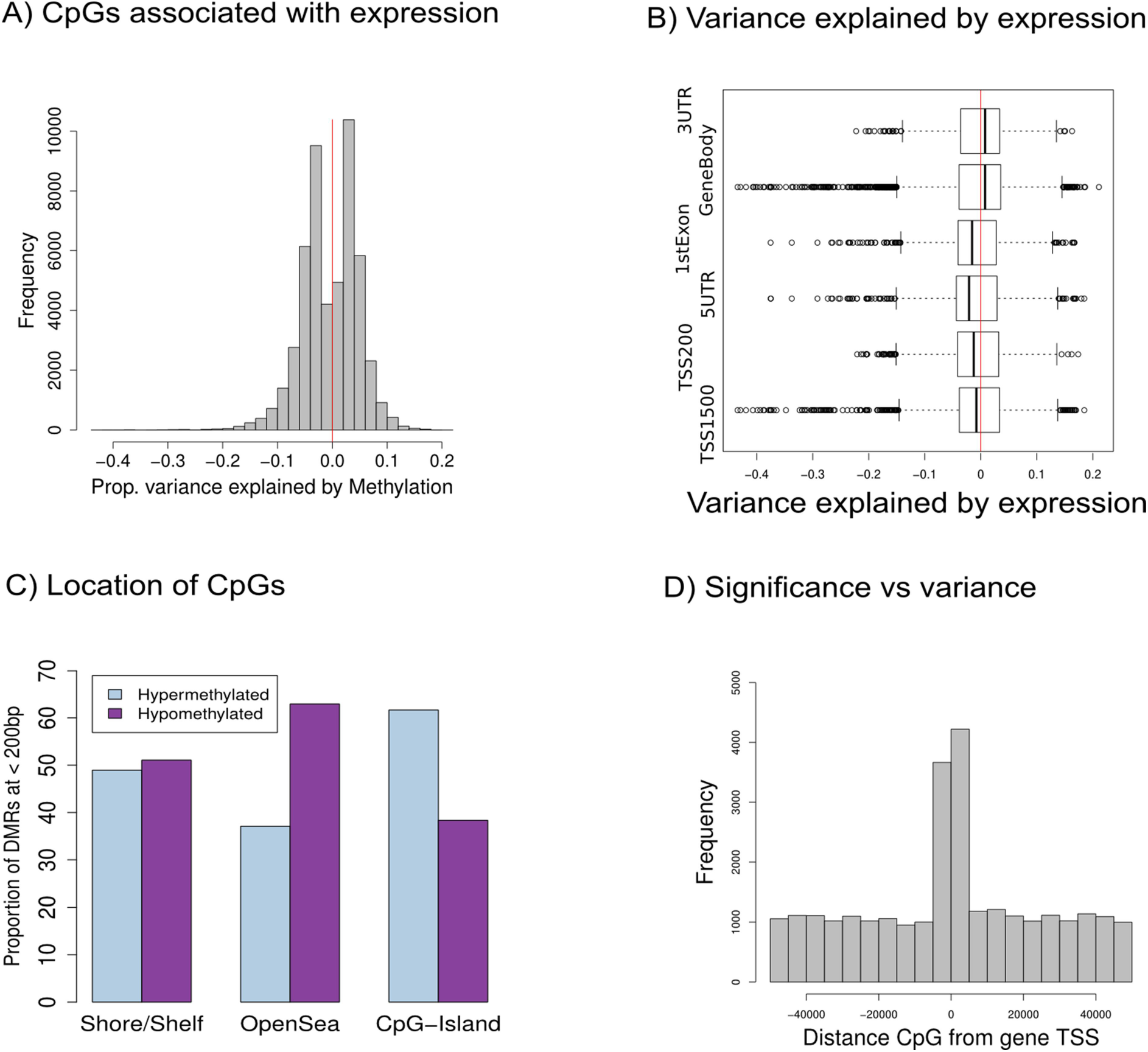
CpGs islands associated to expression around the TSS of the genes. A) Histogram showing the frequency of methylation markers based on the amount of variance explained in expression by methylation levels of nearby CpGs. The proportion of hyper-and hypomethylated CpGs associated to expression levels is near to 50%, with more stronger effects observed for hypomethylated CpGs associated to expression. B) Boxplots showing the genomic position of CpGs associated with expression. C) Barplots showing the percentage of CpGs associated to expression located in shores, open seas and CpG-Island regions. D) Histogram showing the distance of the methylation markers from the TSS of the genes they are associated to. In general, we observed a slightly larger number of methylation marker associated to expression in the promotor and coding region of the genes.

**Supp Figure S8.**
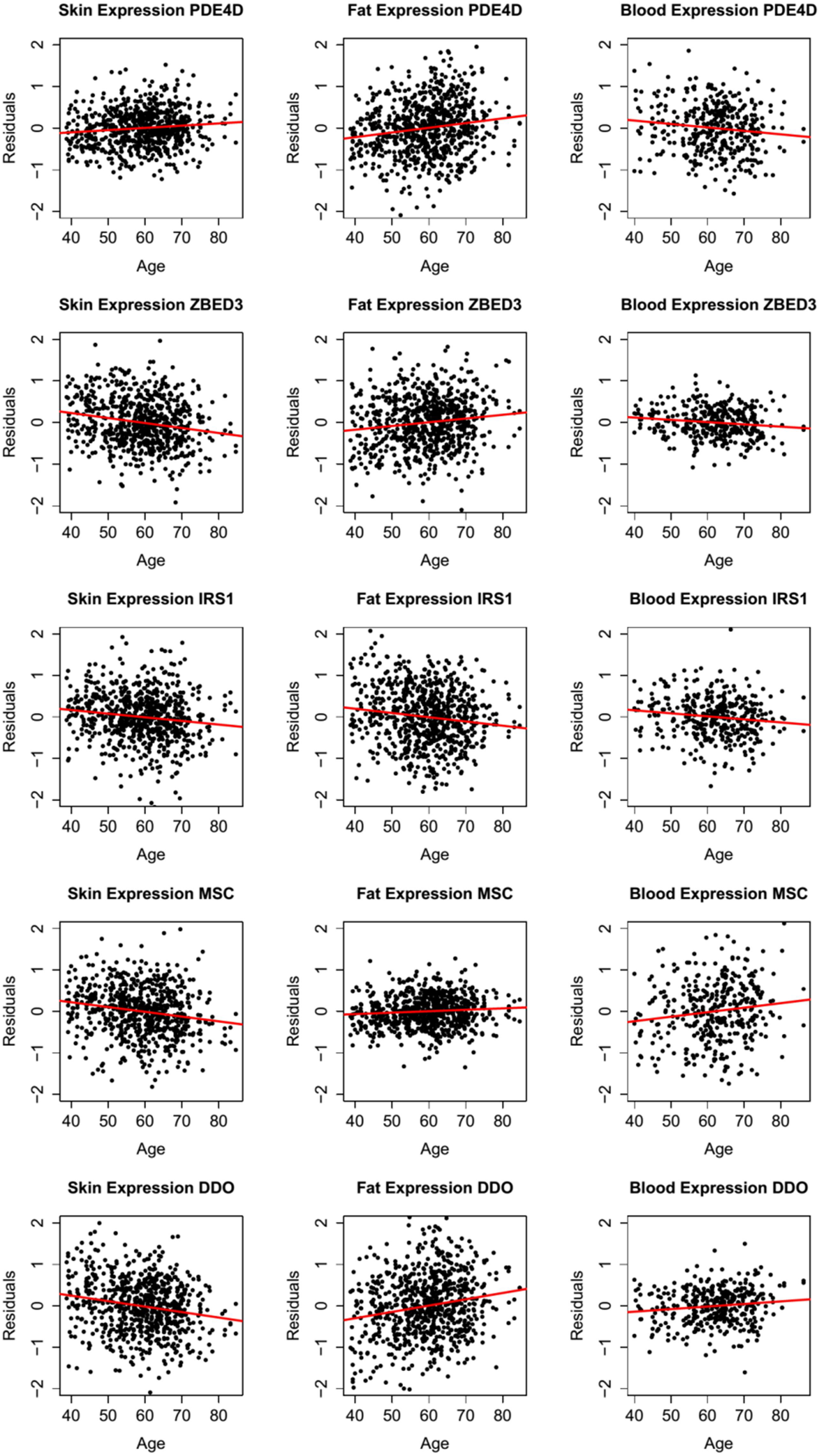
Genes commonly affected by age in the three primary tissues: skin, fat and whole blood. The graph shows the residuals from a linear mixed model removing technical covariates and family structure. The red lines are linear models fitted with the residuals association with age, indicating therefore the direction of effect of age in the expression of the exons plotted.

**Additional Figure 9.**
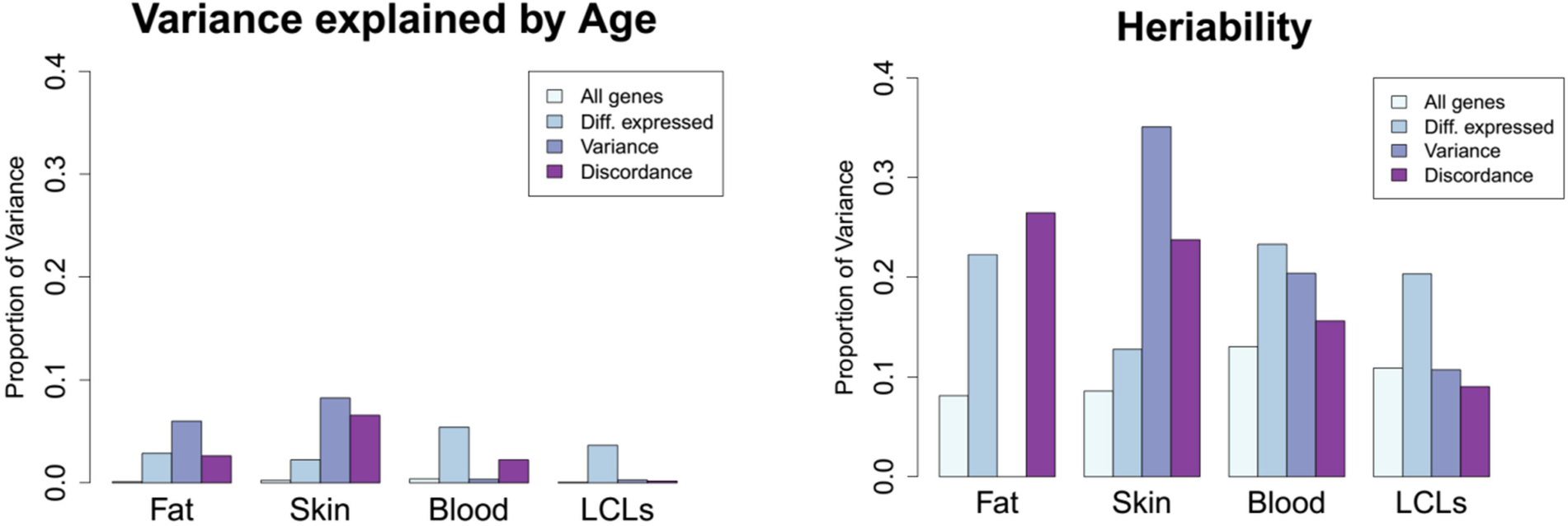
Median proportion of variance explained by age (left) and genetics (right) in all tested genes (All genes), differentially expressed genes with age (diff. expressed), genes changing variance with age (Variance) and genes discordant in MZ twins with age (Discordant). In general, the amount of variance explained by age and heritability in genes significantly affected by age in different ways is larger than in the median of the whole genome. The exception applies to those groups of genes with very little number of genes, like discordance genes in fat with 1 gene. The complete variance decomposition analysis is shown in table S3.

**Additional Table S1.** Number of monozyigous (MZ), dizygous (DZ) and unrelated individuals (individuals with no relatives in the dataset) included in the final analysis per tissue are described on the following.

|  | Fat | Skin | LCLs | Whole Blood |
| --- | --- | --- | --- | --- |
| **Samples** | 766 | 716 | 814 | 384 |
| **MZ pairs** | 131 | 114 | 137 | 69 |
| **DZ pairs** | 187 | 173 | 217 | 91 |
| **Unrelated** | 130 | 142 | 106 | 64 |

**Additional Table S2.**
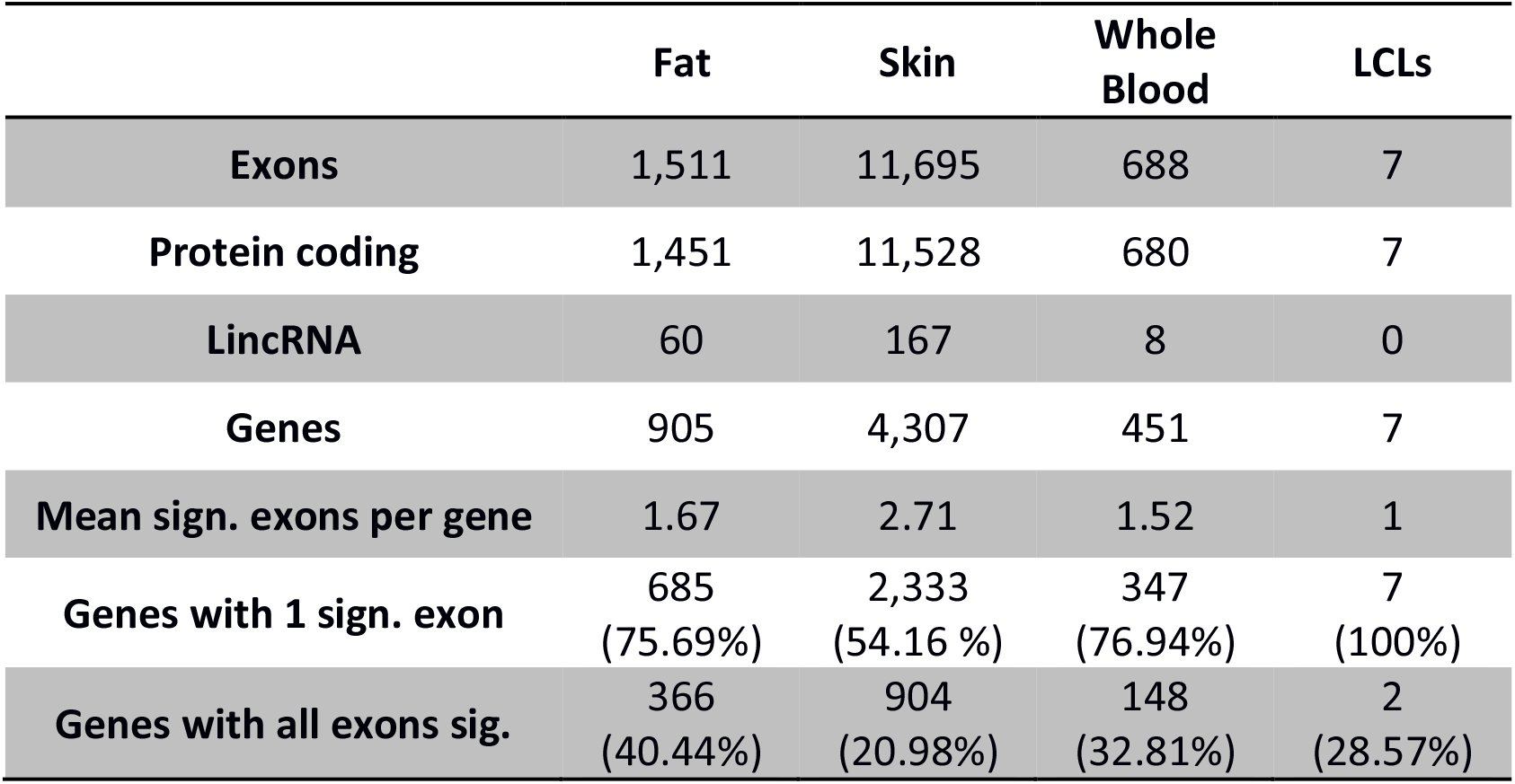
Number of exons and genes significantly associated with age in all tissues. The numbers of those genes that are protein coding genes and LincRNAs are also indicated. The last three rows show the average number of exons per significant gene, number of genes with only exons significantly associated with age and number of genes with all the exons significantly associated with age, respectively. The percentage of the age associated genes are show under each number.

|  | Fat | Skin | Whole Blood | LCLs |
| --- | --- | --- | --- | --- |
| **Exons** | 1,511 | 11,695 | 688 | 7 |
| **Protein coding** | 1,451 | 11,528 | 680 | 7 |
| **LincRNA** | 60 | 167 | 8 | 0 |
| **Genes** | 905 | 4,307 | 451 | 7 |
| **Mean sign. exons per gene** | 1.67 | 2.71 | 1.52 | 1 |
| **Genes with 1 sign. exon** | 685<br>(75.69%) | 2,333<br>(54.16 %) | 347<br>(76.94%) | 7<br>(100%) |
| **Genes with all exons sig.** | 366<br>(40.44%) | 904<br>(20.98%) | 148<br>(32.81%) | 2<br>(28.57%) |

**Additional Table S3.**
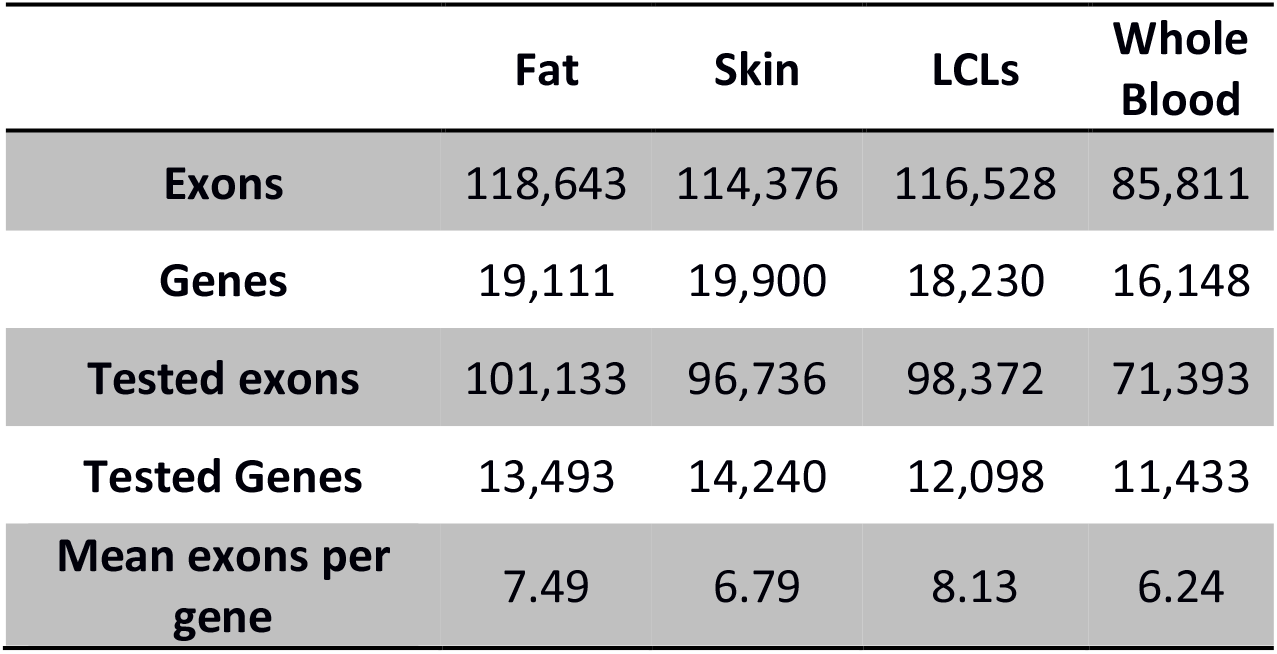
Total number of exons and genes sequenced per tissue, as well as the total number of exons, genes used in the analysis here presented. We use genes defined as protein coding in the GENCODE v10 annotation removing genes with more than 10% zero read count in each tissue. For the analysis presented in this paper, only exons from protein coding genes and LincRNAs from verified loci (level 1) and manually annotated (level 2) were investigated.

**Additional Table S4.** Total number of links identify per tissue, as well as the total number of links per gene detected is shown in the following table. Those link belong to genes included in table 3.

|  | **Fat** | **Skin** | **LCLs** | **Whole Blood** |
| --- | --- | --- | --- | --- |
| **Links** | 221,057 | 179,012 | 259,903 | 184,105 |
| **Genes** | 4,572 | 4,289 | 5,502 | 5,185 |
| **Tested links** | 179,675 | 146,908 | 227,594 | 86,777 |
| **Tested Genes** | 3,876 | 3,557 | 4,673 | 1,590 |

**Additional Table S5.** List of significant exons differentially expressed with age in all tissues.

| Exon | GeneName | Chr | Strand | TSS | P value Fat | Beta Age Fat | P value Skin | Beta Age Skin | P value Blood | Beta Age Blood |
| --- | --- | --- | --- | --- | --- | --- | --- | --- | --- | --- |
| ENSG00000113448.11_58264865_58270907 | PDE4D | chr5 | - | 59284544 | 2.15E-07 | 0.02078 | 0.00103 | 0.00966 | 0.00074 | -0.01471 |
| ENSG00000132846.5_76367897_76373720 | ZBED3 | chr5 | - | 76375207 | 2.49E-05 | 0.01769 | 3.22E-09 | -0.02289 | 0.00166 | -0.01710 |
| ENSG00000169047.4_227659705_227664475 | IRS1 | chr2 | - | 227657101 | 1.09E-05 | -0.01876 | 0.00015 | -0.01187 | 0.00176 | -0.01918 |
| ENSG00000178860.8_72753784_72754982 | MSC | chr8 | - | 72754982 | 0.00113 | 0.01467 | 1.59E-08 | -0.02374 | 0.00018 | 0.02339 |
| ENSG00000203797.4_110712974_110714545 | DDO | chr6 | - | 110713980 | 1.54E-08 | 0.02357 | 3.72E-08 | -0.02299 | 0.00165 | 0.01926 |

**Additional Table S6.** Summary of mean proportion of variance attributed to age, genetics, common environment and unique environment, for exons affected by age in their variance (Variance) and exons discordant for expression with age (Discordant). The last row indicates the number of exons significant for each category (corrected *P*value < 0.05).

|  | Fat |  |  |  | Skin |  |  |  | Blood |  |  |  | LCLs |  |  |  |
| --- | --- | --- | --- | --- | --- | --- | --- | --- | --- | --- | --- | --- | --- | --- | --- | --- |
|  | All | DE | Variance | Discordance | All | Age | Variance | Discordance | All | Age | Variance | Discordance | All | Age | Variance | Discordance |
| Age | 0.0012 | 0.0287 | 0.0601 | 0.0263 | 0.0026 | 0.0224 | 0.0822 | 0.0658 | 0.0039 | 0.0542 | 0.0035 | 0.0225 | 0.0006 | 0.0366 | 0.0030 | 0.0018 |
| Genetics | 0.0809 | 0.2228 | 8.9e-14 | 0.2649 | 0.0856 | 0.1275 | 0.3513 | 0.2380 | 0.1301 | 0.2333 | 0.2037 | 0.1560 | 0.1089 | 0.2032 | 0.1070 | 0.0901 |
| Commo Env. | 0.0236 | 0.0573 | 0.172 | -0.1066 | 0.0000 | 0.0162 | -0.1017 | -0.066 | 0.0380 | 0.0341 | 0.0716 | 0.0293 | 0.0971 | 0.1312 | 0.1966 | 0.0599 |
| Unique Env. | 0.8550 | 0.6655 | 0.690 | 0.8094 | 0.8566 | 0.7662 | 0.6778 | 0.7512 | 0.7403 | 0.6214 | 0.6055 | 0.7179 | 0.7320 | 0.6214 | 0.6224 | 0.7644 |
| N. of exons | 101,133 | 1,511 | 3 | 1 | 96,736 | 11,695 | 239 | 40 | 71,393 | 688 | 13 | 5 | 98,372 | 7 | 3 | 2 |

## References

1. Valdes AM, Glass D, Spector TD: Omics technologies and the study of human ageing. Nat Rev Genet 2013, 14:601–607.

2. López-Otín C, Blasco MA, Partridge L, Serrano M, Kroemer G: The Hallmarks of Aging. Cell, 153:1194–1217.

3. Glass D, Vinuela A, Davies M, Ramasamy A, Parts L, Knowles D, Brown A, Hedman A, Small K, Buil A, et al: Gene expression changes with age in skin, adipose tissue, blood and brain. Genome Biology 2013, 14:R75.

4. Rodwell GEJ, Sonu R, Zahn JM, Lund J, Wilhelmy J, Wang L, Xiao W, Mindrinos M, Crane E, Segal E, et al: A Transcriptional Profile of Aging in the Human Kidney. PLoS Biol 2004, 2:e427.

5. de Magalhaes JP, Curado J, Church GM: Meta-analysis of age-related gene expression profiles identifies common signatures of aging. Bioinformatics 2009, 25:875–881.

6. Viñuela A, Snoek LB, Riksen JAG, Kammenga JE: Genome-wide gene expression regulation as a function of genotype and age in C. elegans. Genome Research 2010, 20:929–937.

7. Li Z, Wright FA, Royland J: Age-Dependent Variability in Gene Expression in Male Fischer 344 Rat Retina. Toxicological Sciences 2009, 107:281–292.

8. Yang J, Huang T, Petralia F, Long Q, Zhang B, Argmann C, Zhao Y, Mobbs CV, Schadt EE, Zhu J, Tu Z: Synchronized age-related gene expression changes across multiple tissues in human and the link to complex diseases. Scientific Reports 2015, 5:15145.

9. Peters MJ, Joehanes R, Pilling LC, Schurmann C, Conneely KN, Powell J, Reinmaa E, Sutphin GL, Zhernakova A, Schramm K, et al: The transcriptional landscape of age in human peripheral blood. Nat Commun 2015, 6.

10. Lu T, Pan Y, Kao S-Y, Li C, Kohane I, Chan J, Yankner BA: Gene regulation and DNA damage in the ageing human brain. Nature 2004, 429:883–891.

11. McCarroll SA, Murphy CT, Zou S, Pletcher SD, Chin CS, Jan YN, Kenyon C, Bargmann CI, Li H: Comparing genomic expression patterns across species identifies shared transcriptional profile in aging. Nat Genet 2004, 36:197 – 204.

12. Bahar R, Hartmann CH, Rodriguez KA, Denny AD, Busuttil RA, Dolle MET, Calder RB, Chisholm GB, Pollock BH, Klein CA, Vijg J: Increased cell-to-cell variation in gene expression in ageing mouse heart. Nature 2006, 441:1011–1014.

13. Somel M, Khaitovich P, Bahn S, Pääbo S, Lachmann M: Gene expression becomes heterogeneous with age. Current Biology 2006, 16:R359–R360.

14. Brinkmeyer-Langford CL, Guan J, Ji G, Cai JJ: Aging Shapes the Population-Mean and -Dispersion of Gene Expression in Human Brains. Frontiers in Aging Neuroscience 2016, 8.

15. Brown AA, Buil A, Viñuela A, Lappalainen T, Zheng H-F, Richards JB, Small KS, Spector TD, Dermitzakis ET, Durbin R: Genetic interactions affecting human gene expression identified by variance association mapping. 2014.

16. Buil A, Brown AA, Lappalainen T, Vinuela A, Davies MN, Zheng H-F, Richards JB, Glass D, Small KS, Durbin R, et al: Gene-gene and gene-environment interactions detected by transcriptome sequence analysis in twins. Nat Genet 2015, 47:88–91.

17. Lappalainen T, Sammeth M, Friedlander MR, t Hoen PAC, Monlong J, Rivas MA, Gonzalez-Porta M, Kurbatova N, Griebel T, Ferreira PG, et al: Transcriptome and genome sequencing uncovers functional variation in humans. Nature 2013, 501:506–511.

18. Horvath S: DNA methylation age of human tissues and cell types. Genome Biology 2013, 14:R115.

19. Tollervey JR, Wang Z, Hortobágyi T, Witten JT, Zarnack K, Kayikci M, Clark TA, Schweitzer AC, Rot G, Curk T, et al: Analysis of alternative splicing associated with aging and neurodegeneration in the human brain. Genome Research 2011, 21:1572–1582.

20. Mazin P, Xiong J, Liu X, Yan Z, Zhang X, Li M, He L, Somel M, Yuan Y, Phoebe Chen YP, et al: Widespread splicing changes in human brain development and aging. 2013.

21. Harries LW, Hernandez D, Henley W, Wood AR, Holly AC, Bradley-Smith RM, Yaghootkar H, Dutta A, Murray A, Frayling TM, et al: Human aging is characterized by focused changes in gene expression and deregulation of alternative splicing. Aging Cell 2011, 10:868–878.

22. Ongen H, Dermitzakis Emmanouil T: Alternative Splicing QTLs in European and African Populations. The American Journal of Human Genetics 2015, 97:567–575.

23. Paré G, Cook NR, Ridker PM, Chasman DI: On the Use of Variance per Genotype as a Tool to Identify Quantitative Trait Interaction Effects: A Report from the Women's Genome Health Study. PLoS Genet 2010, 6:e1000981.

24. Kent JW, Göring HHH, Charlesworth JC, Drigalenko E, Diego VP, Curran JE, Johnson MP, Dyer TD, Cole SA, Jowett JBM, et al: Genotype × age interaction in human transcriptional ageing. Mechanisms of ageing and development 2012, 133:581–590.

25. Yao C, Joehanes R, Johnson AD, Huan T, Esko T, Ying S, Freedman JE, Murabito J, Lunetta KL, Metspalu A, et al: Sex- and age-interacting eQTLs in human complex diseases. Human Molecular Genetics 2014, 23:1947–1956.

26. Wheeler HE, Kim SK: Genetics and genomics of human ageing. Philosophical Transactions of the Royal Society B: Biological Sciences 2011, 366:43–50.

27. Bell JT, Tsai P-C, Yang T-P, Pidsley R, Nisbet J, Glass D, Mangino M, Zhai G, Zhang F, Valdes A, et al: Epigenome-Wide Scans Identify Differentially Methylated Regions for Age and Age-Related Phenotypes in a Healthy Ageing Population. PLoS Genet 2012, 8:e1002629.

28. Hannum G, Guinney J, Zhao L, Zhang L, Hughes G, Sadda S, Klotzle B, Bibikova M, Fan J-B, Gao Y, et al: Genome-wide Methylation Profiles Reveal Quantitative Views of Human Aging Rates. Molecular cell 2013, 49:359–367.

29. Melé M, Ferreira PG, Reverter F, DeLuca DS, Monlong J, Sammeth M, Young TR, Goldmann JM, Pervouchine DD, Sullivan TJ, et al: The human transcriptome across tissues and individuals. Science 2015, 348:660–665.

30. Jeck WR, Siebold AP, Sharpless NE: Review: a meta-analysis of GWAS and age-associated diseases. Aging Cell 2012, 11:727–731.

31. Beekman M, Blanché H, Perola M, Hervonen A, Bezrukov V, Sikora E, Flachsbart F, Christiansen L, De Craen AJM, Kirkwood TBL, et al: Genome-wide linkage analysis for human longevity: Genetics of Healthy Aging Study. Aging Cell 2013, 12:184–193.

32. Chasman DI, Paré G, Mora S, Hopewell JC, Peloso G, Clarke R, Cupples LA, Hamsten A, Kathiresan S, Mälarstig A, et al: Forty-Three Loci Associated with Plasma Lipoprotein Size, Concentration, and Cholesterol Content in Genome-Wide Analysis. PLoS Genet 2009, 5:e1000730.

33. Rodriguez S, Coppede F, Sagelius H, Eriksson M: Increased expression of the Hutchinson-Gilford progeria syndrome truncated lamin A transcript during cell aging. Eur J Hum Genet 2009, 17:928–937.

34. McClintock D, Ratner D, Lokuge M, Owens DM, Gordon LB, Collins FS, Djabali K: The Mutant Form of Lamin A that Causes Hutchinson-Gilford Progeria Is a Biomarker of Cellular Aging in Human Skin. PLoS ONE 2007, 2:e1269.

35. Conneely KN, Capell BC, Erdos MR, Sebastiani P, Solovieff N, Swift AJ, Baldwin CT, Budagov T, Barzilai N, Atzmon G, et al: Human longevity and common variations in the LMNA gene: a meta-analysis. Aging Cell 2012, 11:475–481.

36. Sebastiani P, Solovieff N, DeWan AT, Walsh KM, Puca A, Hartley SW, Melista E, Andersen S, Dworkis DA, Wilk JB, et al: Genetic Signatures of Exceptional Longevity in Humans. PLoS ONE 2012, 7:e29848.

37. Pickrell JK, Pai AA, Gilad Y, Pritchard JK: Noisy Splicing Drives mRNA Isoform Diversity in Human Cells. PLoS Genet 2010, 6:e1001236.

38. Brown A, Ding Z, Viñuela A, Glass D, Parts L, Spector T, Winn J, Durbin R: Pathway Based Factor Analysis of Gene Expression Data Produces Highly Heritable Phenotypes that Associate with Age. G3: GeneslGenomeslGenetics 2015.

39. Gentles AJ, Newman AM, Liu CL, Bratman SV, Feng W, Kim D, Nair VS, Xu Y, Khuong A, Hoang CD, et al: The prognostic landscape of genes and infiltrating immune cells across human cancers. Nat Med 2015, 21:938–945.

40. Nilsson E, Jansson PA, Perfilyev A, Volkov P, Pedersen M, Svensson MK, Poulsen P, Ribel-Madsen R, Pedersen NL, Almgren P, et al: Altered DNA Methylation and Differential Expression of Genes Influencing Metabolism and Inflammation in Adipose Tissue From Subjects With Type 2 Diabetes. Diabetes 2014, 63:2962–2976.

41. Reynolds LM, Taylor JR, Ding J, Lohman K, Johnson C, Siscovick D, Burke G, Post W, Shea S, Jacobs DR Jr, et al: Age-related variations in the methylome associated with gene expression in human monocytes and T cells. Nature Communications 2014, 5:5366.

42. Grundberg E, Small KS, Hedman AK, Nica AC, Buil A, Keildson S, Bell JT, Yang T-P, Meduri E, Barrett A, et al: Mapping cis- and trans-regulatory effects across multiple tissues in twins. Nat Genet 2012, 44:1084–1089.

43. Spector TD, Williams FMK: The UK Adult Twin Registry (TwinsUK). Twin Research and Human Genetics 2006, 9:899–906.

44. Initial sequencing and analysis of the human genome. Nature 2001, 409:860–921.

45. Li H, Durbin R: Fast and accurate short read alignment with Burrows-Wheeler transform. Bioinformatics 2009, 25:1754–1760.

46. Harrow J, Frankish A, Gonzalez JM, Tapanari E, Diekhans M, Kokocinski F, Aken BL, Barrell D, Zadissa A, Searle S, et al: GENCODE: The reference human genome annotation for The ENCODE Project. Genome Research 2012, 22:1760–1774.

47. An integrated map of genetic variation from 1,092 human genomes. Nature 2012, 491:56–65.

48. Howie BN, Donnelly P, Marchini J: A Flexible and Accurate Genotype Imputation Method for the Next Generation of Genome-Wide Association Studies. PLoS Genet 2009, 5:e1000529.

49. Ongen H, Dermitzakis ET: Alternative splicing QTLs in European and African populations using Altrans, a novel method for splice junction quantification. 2015.

50. The R Project for Statistical Computing [http://www.r-project.org/]

51. Bates D, Maechler M, Bolker B: lme4: Linear mixed-effects models using S4 classes. R package version 0.999375-41. 2011.

52. Brown AA: largeQvalue: A program for calculating FDR estimates with large datasets. 2014.

53. Dabney A, Storey JD: qvalue: Q-value estimation for false discovery rate control. R package version 1400.

54. Grundberg E, Meduri E, Sandling Johanna K, Hedman Âsa K, Keildson S, Buil A, Busche S, Yuan W, Nisbet J, Sekowska M, et al: Global Analysis of DNA Methylation Variation in Adipose Tissue from Twins Reveals Links to Disease-Associated Variants in Distal Regulatory Elements. The American Journal of Human Genetics, 93:876–890.

55. Teschendorff AE, Marabita F, Lechner M, Bartlett T, Tegner J, Gomez-Cabrero D, Beck S: A beta-mixture quantile normalization method for correcting probe design bias in Illumina Infinium 450 k DNA methylation data. Bioinformatics 2013, 29:189–196.

56. Visscher PM, Benyamin B, White I: The Use of Linear Mixed Models to Estimate Variance Components from Data on Twin Pairs by Maximum Likelihood. Twin Research (2000) 2004, 7:670–674.

57. Gerrits A, Li Y, Tesson BM, Bystrykh LV, Weersing E, Ausema A, Dontje B, Wang X, Breitling R, Jansen RC, de Haan G: Expression Quantitative Trait Loci Are Highly Sensitive to Cellular Differentiation State. PLoS Genet 2009, 5:e1000692.

58. Anderson M, Braak CT: Permutation tests for multi-factorial analysis of variance. Journal of Statistical Computation and Simulation 2003, 73:85–113.

